# Conservation and evolution of the ribosome across the photosynthetic lineage

**DOI:** 10.64898/2026.07.03.736309

**Authors:** David Pearce, Noah N. Turner, William Tasker-Brown, Sofie van Dorst, Ishika Pramanick, Vinod K. Vogirala, Aled T. Griffith, Gaia Galiberti, Andrew R. J. Curson, Benoît Michel, Jake Richardson, Saima Rehman, Carlo Martins, Gerhard Saalbach, Florent Waltz, David J. Lea-Smith, Benjamin D. Engel, Michael W. Webster

## Abstract

Regulated biogenesis of photosynthetic proteins by the chloroplast ribosomes is central to plant development and environmental adaptation. Here, we describe changes to the ribosome that occurred during evolution of chloroplasts from cyanobacteria in the photosynthetic lineage based on structural, proteomic and bioinformatic analyses. We identify structural features common to oxygenic photosynthetic organisms and distinct from other bacteria, as well as the gain and loss of ribosomal proteins, rRNA modifications and hibernation factors upon endosymbiosis. We uncover structural variation among land plant lineages at sites that remodel the peptide exit tunnel and features associated with translation initiation. The data challenge the prevailing view of ribosomal features that are specific to the chloroplast by demonstrating that many originated earlier, in cyanobacteria, or later, during land plant evolution.

## Introduction

The chloroplast ribosome synthesizes the approximately 85 plant proteins encoded by the chloroplast genome. This includes proteins that perform photosynthesis or support their expression, such as components of the ribosome and RNA polymerase. The products of chloroplast translation comprise most of the leaf protein biomass, and a substantial fraction of available energy is dedicated to the biogenesis and activity of the chloroplast ribosome (1). The chloroplast ribosome resembles that of bacteria due to its inheritance during the endosymbiotic event that produced the chloroplast from a cyanobacterial ancestor. The extent of similarity is notable given dramatic changes in the plastid genome and its gene expression regulatory pathways that have occurred in the approximately 2 billion years since endosymbiosis (2, 3).

Proteomic analysis of the chloroplast ribosome of *Spinacia oleracea* (spinach hereafter) identified homologs of 52 of the 54 *Escherichia coli* ribosomal proteins, five additional proteins absent from *E. coli* termed ‘plastid-specific ribosomal proteins’ (PSRPs 2-6), and a homolog of hibernation promoting factor (HPF) named PSRP1 (4-7). Chloroplast ribosomal proteins are generally larger than their bacterial counterparts due to plastid-specific terminal extensions. The possible roles of PSRPs and terminal extensions were clarified by structures of the spinach chloroplast ribosome in non-translating states (8-12) and of the chloroplast ribosome of the green alga *Chlamydomonas reinhardtii* (*Chlamydomonas* hereafter) (13). The PSRPs range in importance for development of *Arabidopsis thaliana*, with some essential for growth and others largely dispensable (14). Molecular processes specific to cyanobacteria and chloroplasts, such as thylakoid targeting, may be supported by structural features of the chloroplast ribosome distinct from *E. coli* (10, 12). However, this remains unresolved due to a lack of structural information on the cyanobacterial ribosome.

To better understand how ribosome structural features support photosynthetic protein synthesis and the organellar gene regulatory pathways, we sought to establish those that are unique to the chloroplast and conserved among chloroplast-containing organisms. We present structural models of ribosomes from the cyanobacterium *Synechocystis* sp. PCC 6803 (*Synechocystis* hereafter) and the chloroplast of the land plant *Sinapis alba* (*Sinapis* hereafter) from cryo-EM reconstructions resolved to approximately 2Å. Analysis of ribosome structures in translating and non-translating states alongside proteomic, protein bioinformatic and cryo-ET data provides a new framework for understanding translation in photosynthetic organisms.

## Results

### Structure and composition of the chloroplast and cyanobacterial ribosomes

Ribosomes were isolated from chloroplasts of *Sinapis* and *Synechocystis* by a single-step affinity approach (15). *Sinapis* is a member of the *Brassicaceae* family of plants that includes the model land plant *Arabidopsis thaliana*. Monosomes and polysomes were observed in negatively stained samples, indicating both translating and non-translating states were present (Figure S1A). Consensus cryo-EM reconstructions of the *Sinapis* chloroplast ribosome and the *Synechocystis* ribosome were resolved to approximately 1.8Å and 2.0Å respectively (Figure 1A,B, Supplementary Figures S1,S2, Supplementary Table S1). The resolution of surface regions was substantially improved by optimization of sample preparation and data processing approaches, revealing features of the chloroplast ribosome insufficiently resolved in previous studies for confident interpretation (Methods, Figure S1C-G).

**Figure 1:**
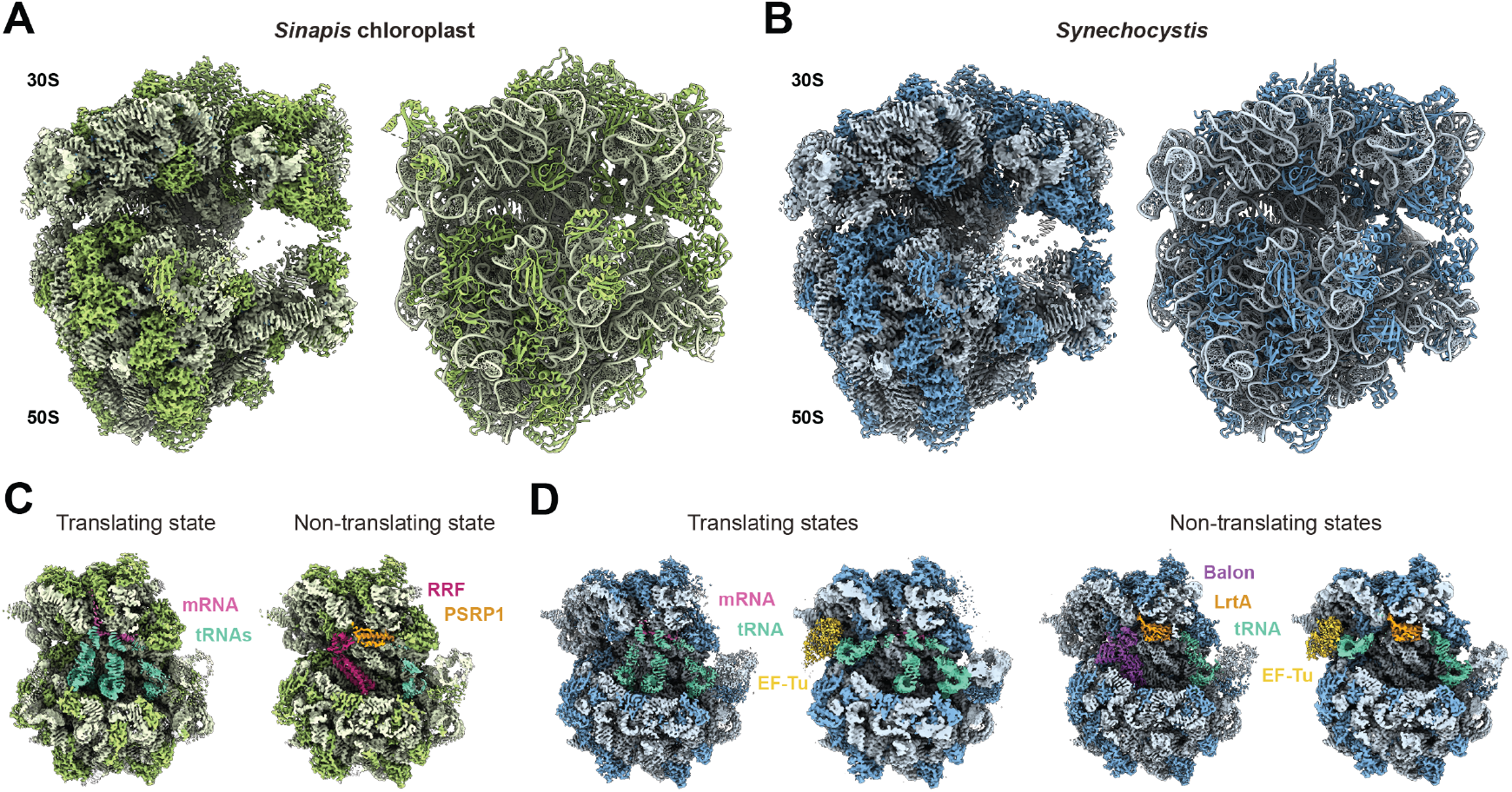
Structures of ribosomes from plant chloroplast and cyanobacteria. **(A**,**B)** Cryo-EM reconstructions (left) and structures (right) of ribosomes from the Sinapis chloroplast and Synechocystis. **(C, D)** Cryo-EM reconstructions of translating and non-translating states of ribosomes from the Sinapis chloroplast and Synechocystis colored by associated factors.

Structural models were generated with ribosomal proteins identified by liquid chromatography-tandem mass spectrometry (LC-MS/MS) (Figure 1A,B, Supplementary Tables S2-S4). We identified *Synechocystis* contains homologs of 53 of the 54 *E. coli* ribosomal proteins and two ribosomal proteins absent from *E. coli*. The first additional protein is Ycf65, a homolog of chloroplast cS23/PSRP3. We rename these bS23 in bacteria and bS23c in chloroplasts for consistent nomenclature (7, 16, 17). The second is an uncharacterized protein that we name bS24. bS24 is absent from the chloroplast ribosome but present in diverse cyanobacteria, suggesting it was lost during endosymbiogenesis (Supplementary Figure S3A).

The only *E. coli* ribosomal protein absent in *Synechocystis* is bL30. The structural data confirm that the absence of bL30 is a shared characteristic of cyanobacterial and chloroplast ribosomes (18). By contrast, bL25 was present in *Synechocystis* but not *Sinapis*, raising a question of the evolutionary timing of its loss. We identified that bL25 is present in diverse cyanobacteria with the notable exception of early-branching lineages most closely related to plastids (*Gloeomargarita lithophora* and *Synechococcus* sp. JA-2-3B) (Supplementary Figure S3A) (3). The protein composition of the chloroplast ribosome therefore not only reflects its cyanobacterial origin, but the specific cyanobacterial clade from which it arose.

The *Sinapis* chloroplast ribosome has the same set of ribosomal proteins as spinach, comprising homologs of 52 *E. coli* proteins and the five PSRPs absent from *E. coli*. Interestingly, whereas *Arabidopsis* contains bS16c encoded by the nuclear genome (19), *Sinapis* contains the product of the plastid bS16c gene, revealing substitution of gene products relatively recently in evolution (Supplementary Tables S2,S3). Proteomic analysis revealed compositional heterogeneity between chloroplast ribosome molecules due to the incorporation of alternative protein isoforms (Supplementary Table S2,S3), as recently observed in the cytosolic ribosome of the land plant *Nicotiana tabacum* (20). At some locations, the cryo-EM density is consistent with the overlapping presence of amino acids that are different between isoforms, providing further support for fine-scale heterogeneity of the chloroplast ribosome (Supplementary Figure S3B). For each protein with multiple isoforms, we modelled the isoform detected with greatest abundance by LC-MS/MS.

The resolution of the cryo-EM reconstruction also supported modelling of uncharacterized fine structural features. We surveyed the cryo-EM density at each nucleotide position for evidence of rRNA modifications. This confirmed the presence of some modification sites detected in the *Arabidopsis* chloroplast ribosome by sequencing approaches (21-26), provided evidence against the presence of others, and identified additional sites not previously reported (Supplementary Figure S3C, Supplementary Table S5). As detailed below, modification events that were likely gained and lost during endosymbiogenesis were distinguished by comparison to the rRNA modifications observed in the cyanobacterial ribosome. The reconstruction of the chloroplast ribosome also resolved rRNA ‘hidden breaks’, the presence of residues arising from RNA editing events, metal ions, and water molecules (Supplementary Figure S3D-F).

Reconstructions were obtained from particle subsets of the *Sinapis* chloroplast ribosome in translating states bound to mRNA and three tRNAs (∼2.7Å resolution), and in non-translating states bound to the HPF homolog PSRP1, E-site tRNA and, in one particle subset, ribosome recycling factor (RRF; ∼2.9Å resolution) (Figure 1C, Supplementary Figure S1H-J). Reconstructions were obtained of translating *Synechocystis* ribosomes either with tRNAs in the A-, P- and E-sites (∼2.5Å resolution) or with EF-Tu and tRNAs in the P- and E-sites (∼2.6Å resolution). Reconstructions were obtained of non-translating *Synechocystis* ribosomes bound to the HPF homolog LrtA and E-site tRNA, and either the hibernation factor Balon (∼2.5Å resolution) or EF-Tu bound to tRNA (∼2.1Å resolution) (Figure 1D, Supplementary Figure S2G-H). Structural models were generated for each of the states, of which only the non-translating chloroplast ribosome has previously been described (9-12).

### A conserved interaction between bS23 and bS1 is unique to photosynthetic organisms

We identified that bS23/bS23c is the only ribosomal protein that is present in both cyanobacteria and chloroplasts and absent from all other bacteria (Supplementary Figure S3A). The structures reveal bS23/bS23c interacts with bS1/bS1c in a way that is conserved between *Sinapis* and *Synechocystis* (Figure 2A). Structural comparison with the *Chlamydomonas* chloroplast ribosome (13) revealed an equivalent interaction between bS23c and bS1c occurs in green algae, suggesting the role for bS23/bS23c is conserved among photosynthetic organisms. However, approximately 20 C-terminal residues present in cyanobacterial bS23 and green alga bS23c are absent in land plant homologs (Figure 2A).

**Figure 2:**
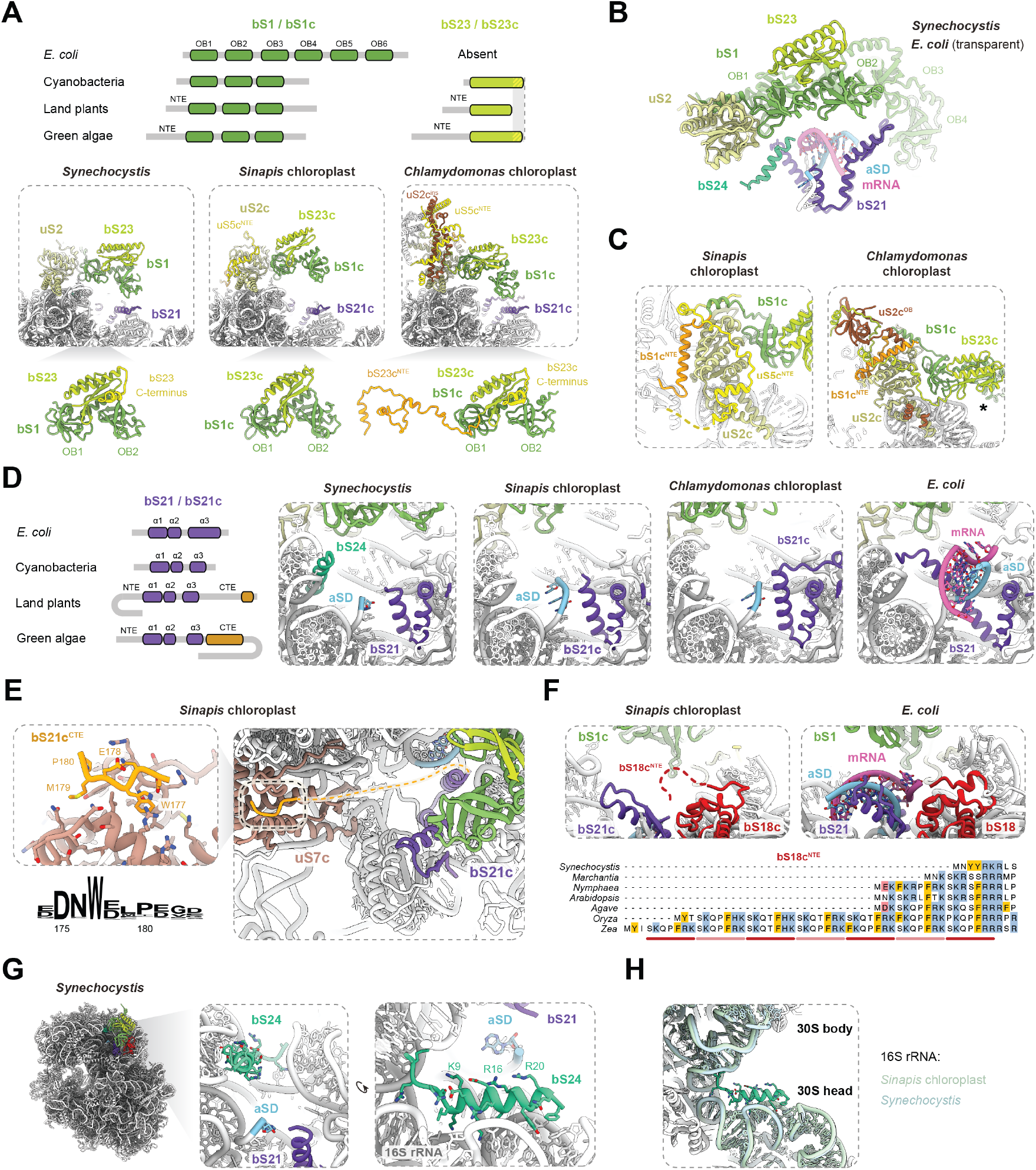
Distinct ribosome features associated with translation initiation in photosynthetic organisms. **(A)** The structural arrangement of bS1 and bS23 in a cyanobacteria (*Synechocystis*) is preserved in chloroplasts of a land plant (*Sinapis*) and a green alga (*Chlamydomonas*). Domain organization diagrams show the presence of conserved domains and N-terminal extensions (NTE) unique to chloroplast proteins. **(B)** Structural overlay of the *Synechocystis* ribosome (opaque) with *E. coli* ribosome in the translation initiation state of mRNA delivery (transparent) shows the position of bS23 relative to the expected position of the mRNA and anti-Shine-Dalgarno (aSD) motif. **(C)** The bS1c N-terminal extension (bS1cNTE) contacts a distinct surface of the chloroplast ribosome in plants (*Sinapis*) and algae (*Chlamydomonas*). The N-terminal extension of uS5c (uS5c^NTE^) and an OB domain insertion in uS2c (uS2c^OB^) also unique to plants and algae respectively are shown. **(D)** Variation in bS21/bS21c position within the 30S platform cavity repositions the aSD in cyanobacteria (*Synechocystis*) and chloroplasts (*Sinapis* and *Chlamydomonas*) relative to *E. coli*. **(E)** Interaction between the bS21c N-terminal extension (bS21c^NTE^) and uS7c in the chloroplast ribosome of land plants. Relative conservation of bS21^NTE^ residues among land plants shows an invariant tryptophan involved in binding. **(F)** The bS18c N-terminal extension (bS18c^NTE^) contains a repeating motif that projects into the 30S platform cavity. Sequences are shown for *Synechocystis, Marchantia polymorpha, Nymphaea colorado, Arabidopsis thaliana, Agave attenuate, Oryza sativa* and *Zea mays*. **(G)** bS24 (hypothetical protein MYO_111350, GenBank: AGF51389) is a conserved cyanobacterial protein located within the 30S platform cavity adjacent to the aSD. **(H)** In *Synechocystis*, bS24 contacts 16S rRNA elements in the 30S head and 30S body domains that are conserved in structure in the chloroplast ribosome (*Sinapis*). In panels A, B, D and F, structural comparisons are made with the *Chlamydomonas* chloroplast ribosome (PDB:9TVU) (13) and *E. coli* ribosome (PDB:9GUT) (29).

bS23/bS23c comprises a single domain that interacts extensively with the helical linker between the OB1 and OB2 domains of bS1/bS1c. Together, they are positioned above the cleft between the 30S head and body domains, equivalent to the first two OB domains of *E. coli* bS1 (27-29).

bS23/bS23c therefore overhangs the mRNA exit channel and the position where the Shine-Dalgarno duplex is bound during initiation (Figure 2B). The conservation of bS23/bS23c residues at the OB1-OB2 domain interface supports a model in which it stabilizes the positions of the OB domains with respect to each other (Supplementary Figure S4A). bS23/bS23c potentially contributes to mRNA binding via a loop containing charged residues proximal to the RNA-binding loops of bS1/bS1c.

The location of bS23/bS23c leads us to speculate that its presence in the photosynthetic lineage is functionally correlated with the distinct domain architecture of bS1c: chloroplast and cyanobacterial homologs of bS1/bS1c have three OB domains, whereas *E. coli* bS1 contains six (Figure 2A) (30). The C-terminal bS1 OB domains of *E. coli* contribute to the positioning of the N-terminal OB domains during initiation and provide additional affinity for mRNA (30), which are roles bS23/bS23c may therefore contribute to in the photosynthetic lineage.

bS23c was previously modelled in ambiguous density in the 30S mRNA exit channel or 30S foot of the spinach chloroplast ribosome (named PSRP3/cS23 in these studies) (10, 11). The improved resolution of our reconstructions allowed us to assign these instead to parts of PSRP1 and cS22/PSRP2 (Supplementary Figure S4B). We resolved bS1c-bound bS23c in cryo-EM reconstructions only when the surfactant CHAPSO was included during cryo-EM sample preparation, suggesting it is prone to damage at the air-water interface like bS1 (Figure S1E). It also supported assignment of density that encircles uS2c to the N-terminal extensions of bS1c (bS1c^NTE^) and uS5c (bS5c^NTE^) (Figure 2C). The approximate position of the bS1/bS1c OB3 domain was identified in reconstructions of translating states, suggesting its position is stabilized by contact with the exiting mRNA (Supplementary Figure S4C).

### Evolution of ribosome features associated with translation initiation in chloroplasts

Structural differences in how the Shine-Dalgarno duplex is contacted may underpin unusual translation initiation region features identified in cyanobacteria (31, 32) and contribute to mRNA selection during translation initiation in chloroplasts (30). During translation initiation in *E. coli*, the Shine-Dalgarno duplex can be positioned between two α-helices of bS21 (29, 30, 33). We observed that in both *Sinapis* and *Synechocystis* the C-terminal α-helix of bS21/bS21c in cyanobacteria is shorter and interacts with a different surface of the 30S platform cavity (Figure 2D). The first three nucleotides of the *Sinapis* anti-Shine-Dalgarno motif were resolved and are contacted by bS21 residue positions different to those that contact the anti-Shine-Dalgarno motif in *E. coli* (Figure 2D, Supplementary Figure S4D). These residues of bS21 are also conserved in diverse cyanobacteria, suggesting mRNA recruitment to ribosomes in chloroplasts and cyanobacteria has a commonality distinct from *E. coli*.

Chloroplasts differ from cyanobacteria in containing a C-terminal extension of bS21c (bS21c^CTE^) not previously resolved. We observed that bS21c^CTE^ contacts uS7c through four residues near the C-terminus that are conserved across land plants (Figure 2D,E). As uS7c is on the opposite side of the 30S cleft to the bS21c α-helices, the intervening 30 unresolved residues of bS21c must span approximately 60Å directly across the channel in which mRNA can enter during initiation and exits during elongation (Figure 2E). bS21c consequently tethers the 30S head and body domains, potentially affecting the kinetics of their relative movements during different phases of translation. The interaction between bS21c^CTE^ and uS7c does not occur in *Chlamydomonas*, and is not predicted for other green algae, indicating any effects of bS21c^CTE^ are specific to chloroplasts of land plants (Supplementary Figure S4E) (13).

The *Sinapis* ribosome reconstruction contains poorly ordered density projecting into the 30S platform cavity in a position consistent with the N-terminal extension of bS18c (bS18c^NTE^), (Supplementary Figure S4F). In *E. coli*, the N-terminus of bS18 becomes ordered when it contacts mRNA during initiation through positively charged residues (29). We identified the bS18c^NTE^ contains a motif of 7 amino acids that includes basic and aromatic residues which has the potential to make additional contacts with incoming mRNA during initiation or exiting mRNA during elongation (Figure 2F). The number of copies of the motif in the bS18c^NTE^ varies across different plant clades: bryophytes (*Marchantia*) contain a single copy, dicots (*Nymphaea, Arabidopsis*) contain two, and monocots (*Agave, Zea, Oryza*) contain between two and seven. We therefore predict the involvement of bS18c in mRNA binding is greater in chloroplasts than bacteria and variable among plant species.

We identified that bS24 is the only ribosomal protein that is absent from the chloroplast ribosome of land plants and green algae but present in the genome of *G. lithophora*, the cyanobacterium most closely related to chloroplasts (Supplementary Figure S3A). bS24 comprises a single N-terminal α-helix that binds the junction of the 30S head and body domains followed by C-terminal residues that are unresolved and unconserved among diverse bacteria (Figure 2G, Supplementary Figure S4G). The architecture of the 16S rRNA is the same in chloroplasts, suggesting bS24 does not remodel its binding site (Figure 2H). Notably, basic residues of bS24 that do not contact rRNA (K9 and R20; Fig 2G) are highly conserved (Supplementary Figure S4G). Their proximity to the anti-Shine-Dalgarno motif suggests that bS24 may contribute to Shine-Dalgarno duplex contact during initiation like bS21. It is possible that the acquisition of ribosomal protein terminal extensions (bS18c^NTE^ and bS21c^CTE^) functionally compensates for the absence of bS24 in chloroplasts due to their similar location.

### Diversification of the ribosome peptide exit tunnel in land plant chloroplasts

We observed that the ribosome region surrounding the peptide exit tunnel is different between cyanobacteria and other bacteria, between chloroplasts and cyanobacteria, and between different clades of land plants (Figure 3A). The peptide exit tunnel of both the *Synechocystis* and chloroplast ribosomes is wider than that of *E. coli* by approximately 10 Å (Figure 3B, Supplementary Figure S5A). The uL24 loop that overhangs the peptide exit tunnel in *E. coli* is retracted to the tunnel edge in the ribosomes of *Synechocystis, Sinapis* and *Chlamydomonas*. This is likely due to the shortening of the loop by one residue and the absence of a proline at the position equivalent to *E. coli* P50 that produces a bend in the loop (Supplementary Figure S5B). We identified that the uL24 loop varies in length and sequence across diverse bacteria, indicating the photosynthetic lineage differs from *E. coli* but not from all other bacteria (Supplementary Figure S5B).

**Figure 3:**
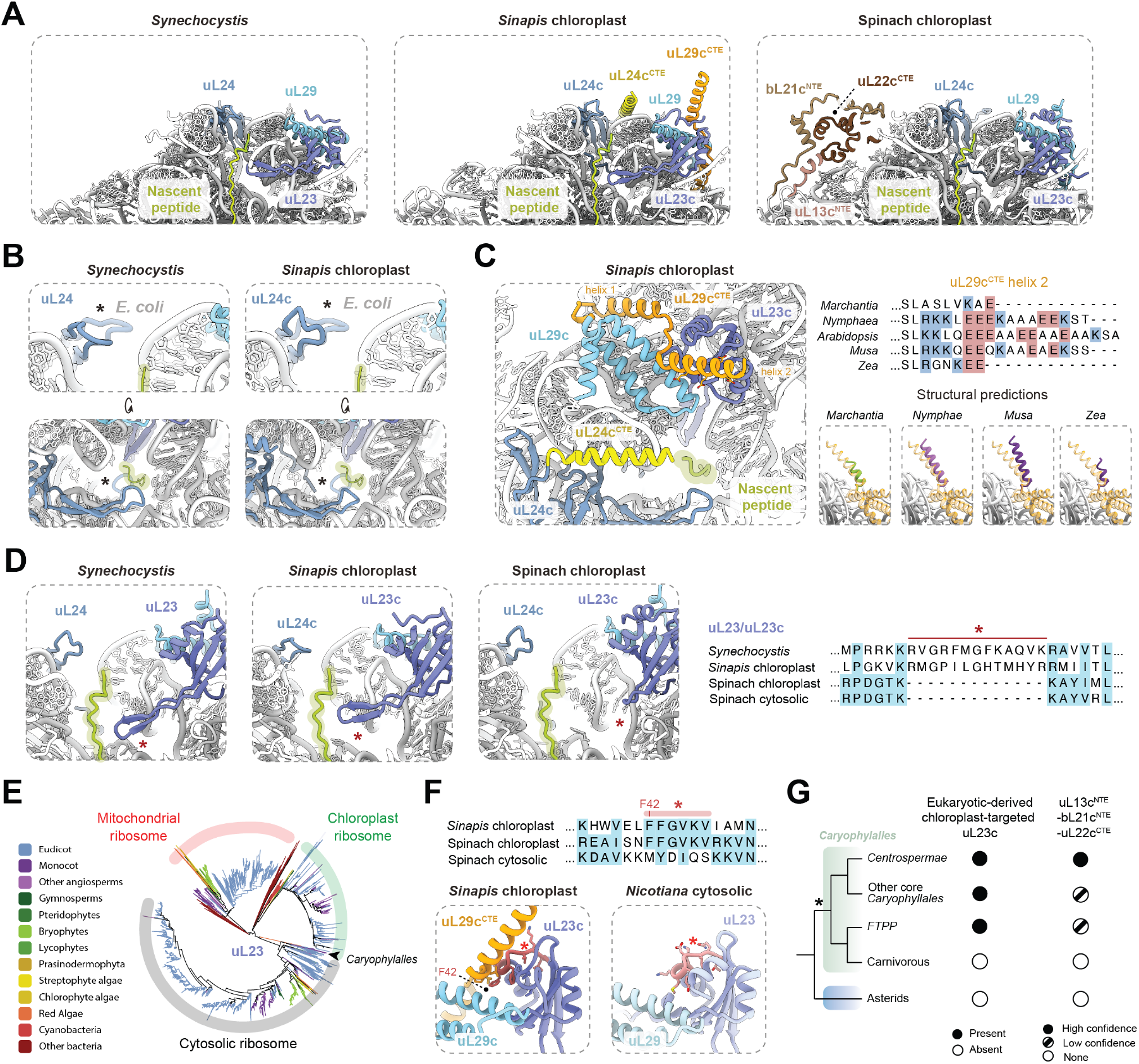
Structural diversity in the chloroplast ribosome peptide exit tunnel. **(A)** Ribosome peptide exit tunnels of *Synechocystis* and chloroplasts of *Sinapis* and spinach (PDB:6ERI) (12). A representative path of nascent peptide is shown (green). **(B)** Structural comparison of the *E. coli* peptide exit tunnel (transparent overlay) with *Synechocystis* (opaque) or the *Sinapis* chloroplast (opaque). The uL24 loop that overhangs the exit tunnel in *E. coli* but not the green lineage is indicated (asterisk). **(C)** C-terminal extensions of uL24c (uL24c^CTE^) and uL29c (uL29c^CTE^) overhang the peptide exit tunnel of the *Sinapis* chloroplast ribosome. The C-terminus of uL29c^CTE^ (‘helix 2’) consistently comprises a charged α-helix that varies in sequence and length across plant species. Structural predictions of the uL29c^CTE^ for representative species are shown overlayed with the *Sinapis* uL29c (orange, transparent). **(D)** The uL23c loop entering the peptide exit tunnel (asterisk) is shorter in spinach than *Sinapis* and *Synechocystis* due to its ancestral similarity to the cytosolic homolog of uL23. **(E)** Phylogenetic analysis of plant uL23 homologs shows sequence similarity of spinach uL23c with homologs in other *Caryophyllales* (arrow), and with uL23 homologs in the cytosolic ribosome. **(F)** Incorporation of eukaryotic-derived uL23c in the chloroplast ribosome of spinach is supported by change in a short motif (red, asterisk) to resemble plastid-encoded uL23c sequences, such as those of *Sinapis*. Structural comparison of the chloroplast ribosome (*Sinapis*) and plant cytosolic ribosome (*Nicotiana;* PDB:8AZW) (20) shows interaction of the motif with uL29c in chloroplasts. **(G)** Correlation of peptide exit tunnel structural features among clades of *Caryophyllales* grouped as in (55).

The ribosomes of plant chloroplasts and cyanobacteria differ in the surface lining the peptide exit tunnel pore due to C-terminal extensions of uL24c (uL24c^CTE^) and uL29c (uL29c^CTE^) in chloroplasts. In *Sinapis*, the α-helices at the termini of uL24c^CTE^ and uL29c^CTE^ overhang the peptide exit tunnel and were resolved upon filtering reconstructions to approximately 4Å resolution (Figure 3C, Supplementary Figure S5C). Analysis of uL29c sequences of diverse plants alongside structural predictions revealed the uL29c^CTE^ consistently comprises a charged α-helix that varies in sequence and length across plant species. In most angiosperms, a series of conserved glutamate residues line one side of the uL29c^CTE^ α-helix, potentially interacting with the emerging nascent chain or factors that engage the ribosome.

A comparison of the sequences of chloroplast ribosomal proteins surrounding the peptide exit tunnel across species revealed that spinach, the model for our current structural understanding of the chloroplast ribosome, contains two distinctive features not shared by most land plants. Firstly, uL23c in the spinach chloroplast is encoded by a nuclear gene that has diverged from the gene encoding the eukaryotic uL23 constituent of the cytosolic ribosome; the diverged spinach uL23c protein has replaced the ancestral prokaryotic uL23c homolog encoded in the chloroplast genome (34). By contrast, *Sinapis* contains uL23c of bacterial origin encoded by a nuclear gene, as does the *Chlamydomonas* chloroplast ribosome (13). Consequently, the loop of uL23/uL23c that enters the peptide exit tunnel in *Sinapis* and *Chlamydomonas* is more like *Synechocystis* and *E. coli*, whereas that of spinach uL23 is 13 residues shorter (Figure 3D, Supplementary Figure S5D). The widened central cavity of the peptide exit tunnel identified in spinach is therefore not a conserved feature that evolved to support thylakoid targeting or provide an alternative exit channel, as was suggested (10, 12).

We identified convergent molecular evolution of features that supported the inclusion of eukaryotic-derived uL23c into the chloroplast ribosome in a common ancestor of the *Caryophyllales* order of superasterids, which includes spinach (Figure 3E,F). Whereas the spinach uL23c sequence largely resembles cytosolic uL23 from which it diverged, a motif of 6 residues instead matches the plastid-encoded uL23c of other plant species (Figure 3F, Supplementary Figure S5E). The motif includes a phenylalanine residue that inserts into a hydrophobic pocket formed by the uL29c^CTE^. Thus, a relatively small but specific sequence change, alongside the acquisition of a chloroplast targeting peptide, was sufficient for the substitution of prokaryotic uL23c for eukaryotic-derived uL23 during land plant evolution.

A second feature of the spinach chloroplast ribosome we identified to be absent from the chloroplast ribosomes of most plant species, and all cyanobacteria, is a domain on the ribosome surface adjacent to the peptide exit tunnel formed by sequence extensions of uL13c, bL21c, uL22c (uL13c^NTE^, bL21c^NTE^, uL22c^CTE^; Figure 3A). This domain expands the surface of the ribosome facing thylakoid proteins during targeting of nascent peptides. We identified that this feature is also unique to *Caryophyllales*, and is largely conserved among its members, revealing a phylogenetic correlation with the presence of eukaryotic-derived uL23c (Figure 3G). The presence of uL13c^NTE^, bL21c^NTE^ and uL22c^CTE^ may compensate for a structural change produced by the incorporation of eukaryotic-derived uL23c (12), but the basis remains unclear from the structural models.

### Origin and function of ribosomal proteins unique to the chloroplast

The roles of PSRPs are clarified by insight into the timing of their evolutionary emergence and correlated changes in structural features surrounding their binding sites. cL38 (also called PSPR6) is the only ribosomal protein present in all examined plants and algae and absent in cyanobacteria, suggesting it could have emerged during endosymbiogenesis (Figure 4A). cL38 was proposed to have evolved to compensate for plastid-specific changes to 5S rRNA architecture based on a comparison to *E. coli* (10). However, the cleft between the 23S and 5S rRNA regions occupied by cL38 is structurally identical in *Synechocystis*, indicating cL38 is not required for rRNA architecture (Supplementary Figure S6A).

**Figure 4:**
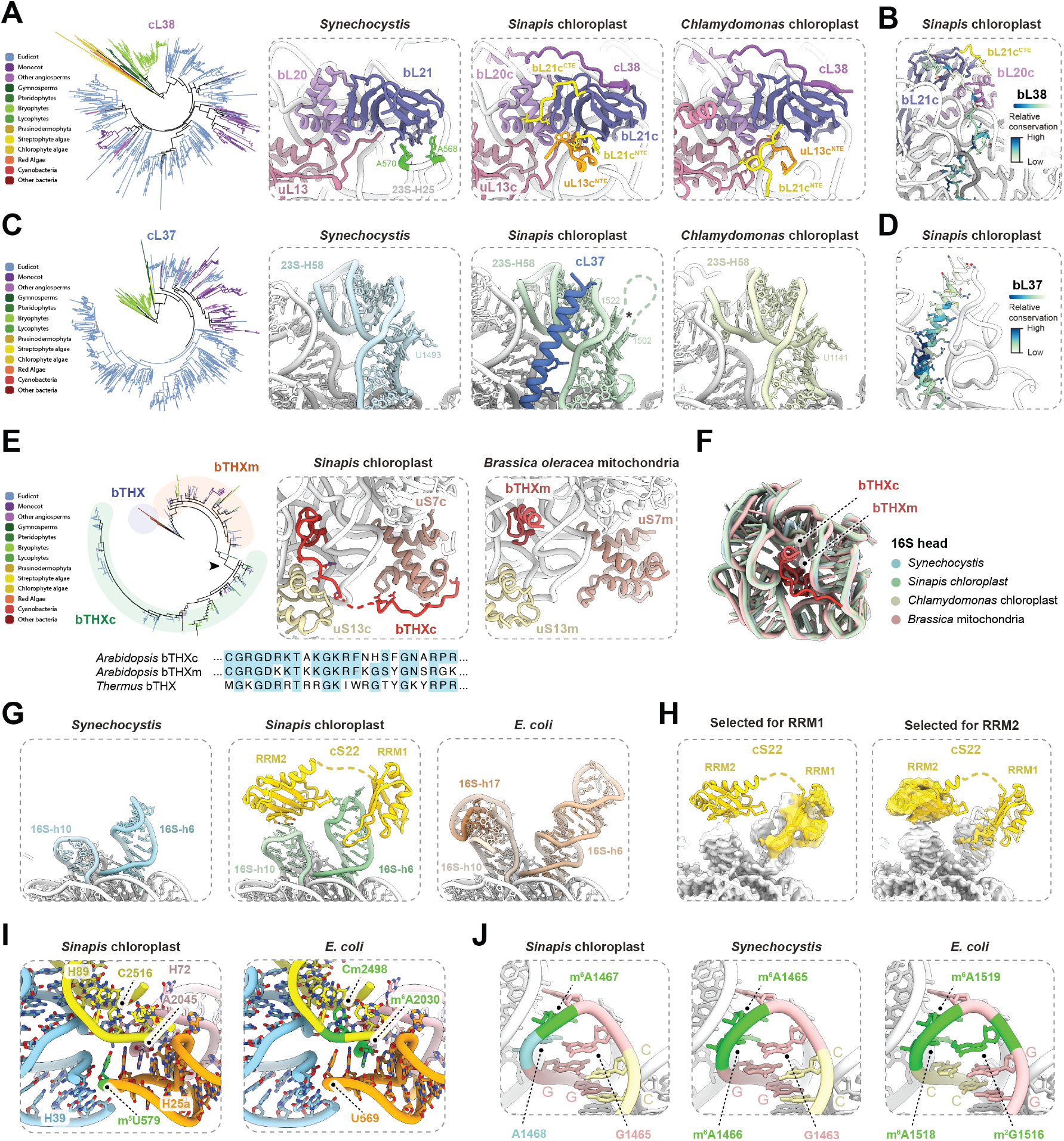
Ribosomal proteins and rRNA modifications unique to the chloroplast ribosome. **(A)** Phylogenetic search for cL38 homologs among plants, algae and bacteria shows conservation among algae and land plants. Binding of bL21c to the chloroplast ribosome is supported by cL38 and plastid-specific terminal extensions of uL13c (uL13^NTE^) and bL21c (bL21^NTE^ and bL21^CTE^). Interactions of bL21 with 16S nucleotides in *Synechocystis* (green) are absent from the chloroplast ribosome. **(B)** Residue-level conservation of cL38. **(C)** Phylogenetic search for cL37 homologs among plants, algae and bacteria shows conservation among land plants, without representatives identified in algae. The presence of cL37 in the chloroplast ribosome of land plants (*Sinapis*) is correlated with the presence of an unstructured insertion in 23S helix 58 (23S h58). These are absent from ribosomes of cyanobacteria (*Synechocystis*) and a green alga chloroplast (*Chlamydomonas*). **(D)** Residue-level conservation of cL37. **(E)** Phylogenetic analysis of plant and bacterial bTHX homologs shows bTHXc is more similar in sequence to bTHXm than to bacterial bTHX. Structural comparison of the interaction of bTHXc with the chloroplast ribosome and bTHXm with the plant mitochondrial ribosome. **(F)** rRNA structure surrounding bTHXm. The 16S architecture is almost identical in ribosomes containing bTHX homologs (*Sinapis* chloroplast and *Brassica oleracea* mitochondria) and those that do not (*Synechocystis* and *Chlamydomonas* chloroplast). **(G)** Structural basis of the interaction between cS22 and 16S rRNA in land plant chloroplasts (*Sinapis*), showing structural similarity of the 16S region in cyanobacteria (*Synechocystis*) and their structural dissimilarity to *E. coli*. **(H)** Cryo-EM reconstructions following focused refinement and classification for each of the RRM domains of cS22. **(I)** Comparison of the region surrounding chloroplast-specific RNA modification site (*Sinapis* 23S m^5^U579) to *E. coli*, which contains two modifications absent in chloroplasts and cyanobacteria (*E. coli* 23S m^6^A2030 and 23S Cm2498). **(J)** 16S-H45 contains fewer RNA modification sites in *Synechocystis* and *Sinapis* chloroplasts than *E. coli*. Structures shown for *Chlamydomonas* chloroplast ribosome (PDB:9TVU) (13), *Brassica oleracea* mitochondrial ribosome (PDB:9GYT) (61) and *E. coli* ribosome (PDB:8PKL) (62).

We instead propose cL38 evolved to support incorporation of bL21c, which is nuclear-encoded and imported to the chloroplast. *Synechocystis* bL21 contacts the 23S and bL20, as in other bacteria, but also contacts 23S helix 25 (23S-H25) that is absent in *E. coli* (Figure 4A). Chloroplasts lack an equivalent interaction between bL21c and 23S-H25. Interaction with cL38 is one of three features unique to the chloroplast that likely stabilize bL21c binding to the ribosome and potentially compensate for otherwise weakened bL21c association; extensions to uL13c and bL21c provide additional interactions connecting bL21c to the ribosome surface. Consistent with this hypothesis, the C-terminal residues of cL38 that contact bL21 are among its most highly conserved among land plants (Figure 4B).

We identified that cL37 (also called PSRP5) is present in land plants but not algae or cyanobacteria (Figure 4C), consistent with recent analysis of the *Chlamydomonas* chloroplast ribosome (13). Comparison of the *Sinapis, Synechocystis* and *Chlamydomonas* structures supports a model in which cL37 emerged in a common ancestor of extant land plants to compensate for loss of independent structural integrity within 23S helix 58 (23S-H58). At a bend in 23S-H58, land plant chloroplasts have ∼18 more nucleotides than cyanobacteria and ∼19 more than *Chlamydomonas* chloroplasts, which were disordered in cryo-EM reconstructions (Figure 4C). cL37 consists of an α-helix that runs parallel to the opposing rRNA strand, making extensive contacts that are expected to counteract increased mobility of the rRNA stem. In support, residues of cL37 that interact with nucleotides that require structural compensation (23S 1544-1546) are among the most conserved (Figure 4D).

bTHXc (also called PSRP4) is conserved among land plants but absent from cyanobacteria and green algae (13). The evolutionary origin of PSRP4/bTHXc is unclear given it has structural similarity to bTHX, which is present in some bacterial lineages, but was seemingly not inherited alongside the rest of the ribosome from the cyanobacterial ancestor of the chloroplast (7). Analysis of bTHXc sequence conservation revealed similarity to the plant mitochondrial bTHX homolog (bTHXm) that suggests common ancestry (Figure 4E). bTHXc is therefore likely derived from a bTHXm gene duplication event and became chloroplast targeted in a common ancestor of extant land plants. bTHXc contains a C-terminal extension absent from bacteria and mitochondria (bTHXc^CTE^), which contacts uS7c through residues conserved among land plants except for spinach and close relatives (Figure 4E, Supplementary Figure S6B,C). The presence of bTHXc does not evidently affect the architecture of the 16S rRNA to which it binds (Figure 4F). The importance of bTHXc to chloroplast biogenesis in *Arabidopsis* (35) suggests it may promote or stabilize 16S folding in a way that, given its absence, is not needed in green algae or cyanobacteria.

The phylogenetic distribution of cS22 (also called PSRP2) suggests it also emerged in a common ancestor of land plants (7). The relative positions of the first and second RRM domains of cS22 were not previously confidently assigned, and we discovered they bind 16S helices 6 (16S-h6) and 10 (16S-h10) respectively (Figure 4G, Supplementary Figure S4C). cS22 RRM1 contacts the groove of 16S-h10 via an extended loop between the second and third β strands. cS22 RRM2 likely contacts a GUU motif at the tip of 16S-h6, consistent with the preference of other chloroplast ribonucleoproteins for G/U-rich sequences (36, 37). Each domain of cS22 was better resolved in different subsets of the imaged particles, suggesting they can move relatively independent of each other (Figure 4H). 16S-h6, 16S-h10 and neighboring 16S-h17 are structurally unlike their *E. coli* counterparts, leading to the suggestion cS22 may have evolved to assist their folding (9, 10). However, the structures of *Synechocystis* and *Sinapis* chloroplast 16S-h6 and 16S-h10 are structurally similar despite the absence of a cS22 homolog in cyanobacteria. We hypothesize that cS22 may, like bTHXc, promote or stabilize 16S folding in a way specifically required by the chloroplasts of land plants.

### Ribosomal RNA modifications in chloroplast and cyanobacteria ribosomes

16 identified rRNA nucleotide modifications were common to *Synechocystis* and *Sinapis*, suggesting they are ancestral to the photosynthetic lineage (Supplementary Table S5). Three of these are unmodified in *E. coli* (16S C1409 and 23S C1920 and G2553) but modified in ribosomes of other species (25, 38-41), indicating they are not unique to photosynthetic organisms. Conversely, 11 rRNA positions modified in *E. coli* were confirmed to be unmodified in *Synechocystis* and *Sinapis*: 3 in 16S, 8 in 23S.

We identified a single rRNA modification unique to the chloroplast that likely arose following endosymbiogenesis (*Sinapis* 23S m^5^U579). This methylation site corresponds to *E. coli* nucleotide U569 located at the tip of helix 25a (Figure 4E). The region contains several modified nucleotides associated with formation of the structured RNA core of the 50S. We observed that chloroplasts and cyanobacteria lack two of the modifications in this region that are present in *E. coli*: m^6^A2030 and Cm2498 (Figure 4E). The position of chloroplast-specific modification m^5^U579 suggests it may provide an alternative route to rRNA structural stability, connecting helix 25a with helices 72 and 89 that contain the unmodified residues at positions equivalent to *E. coli* m^6^A2030 and Cm2498.

A single observable rRNA modification is specifically absent from the chloroplast: *Sinapis* 16S A1468, corresponding to *Synechocystis* m^6^A1466 and *E. coli* m^6^A1519 in 16S helix 45 (16S-h45) (Figure 4F). At a late stage of *E. coli* 30S biogenesis, m^6^A1519 is introduced alongside m^6^A1518 by KsgA in a processive mechanism (42). Chloroplasts have a homolog of KsgA (43), which we hypothesize lacks the processivity of its prokaryotic and eukaryotic homologs such that it modifies only a single adenosine. The rRNA conformations are nonetheless the same in *Sinapis* and *Synechocystis*, contrasting the substantial changes in rRNA architecture observed when modifications are absent from 1518 and 1519 in *Thermus* (44). The preservation of rRNA conformation could be due to other differences in 16S-h45: *Sinapis* and *Synechocystis* lack methylation at positions corresponding to *E. coli* m^2^G1516, and adjacent base pairs are swapped (Figure 4F). We propose that altered patterns of 16S modification are therefore associated with rRNA sequence differences in the photosynthetic lineage.

### Diversification of the mRNA entrance channel in the chloroplast ribosome

Reconstructions of the *Sinapis* and *Synechocystis* ribosomes in states of translation elongation revealed mRNA and tRNA binding sites and density for the path of the nascent peptide. The mRNA entrance channel of the *Sinapis* chloroplast ribosome is less well defined than that of *Synechocystis* due to the flexibility of a uS4c loop that lines the pore (Figure 5A) (12). However, we identified that this is not a general feature of chloroplast ribosomes as the uS4c loop of *Chlamydomonas* is ordered and overhangs the mRNA entrance channel (Figure 5A). Analysis of uS4c sequences suggest the flexibility is common to land plants because residues contacting the 16S (*Synechocystis* Q34 and H35) are conserved across bacteria and algae but absent from all land plants (Figure 5A).

**Figure 5:**
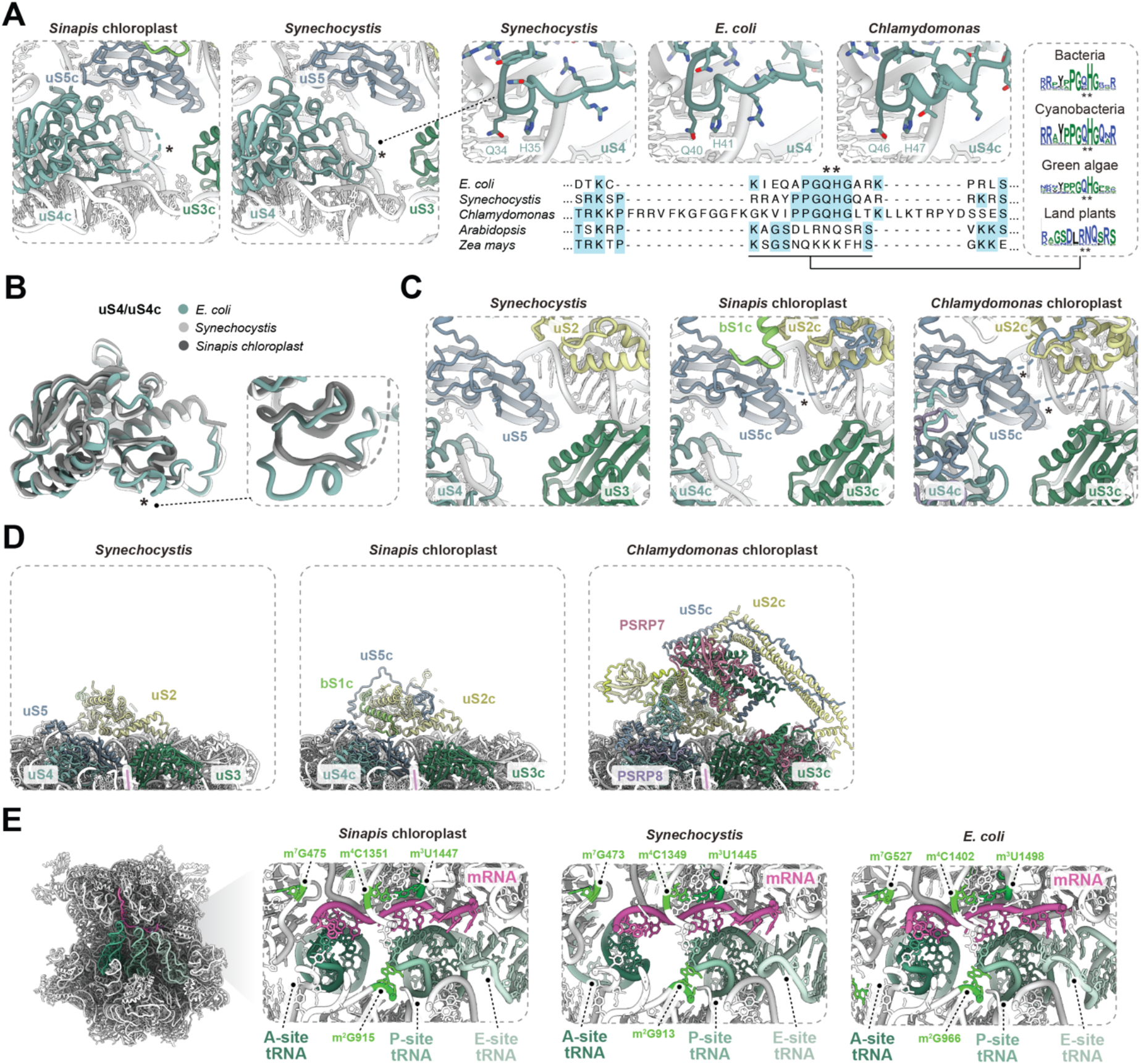
mRNA channel and peptidyl transferase center features in the photosynthetic lineage. **(A)** The uS4/uS4c loop that borders the mRNA entrance channel is not structurally well defined in the ribosome of land plant chloroplasts but is the ribosomes of cyanobacteria (*Synechocystis*) and green algae chloroplasts (*Chlamydomonas*). Comparison of the uS4/uS4c loop in *E. coli*, cyanobacteria (*Synechocystis*) and chloroplasts of green algae (*Chlamydomonas*) reveals conserved contact of 16S by a glutamine and a histidine residue of uS4/uS4c, consistent with analysis of the spinach chloroplast ribosome (12). Analysis of sequence conservation of the uS4/uS4c loop shows the consistent absence of the glutamine and histidine residues in all land plants examined. **(B)** Structural overlay of uS4/uS4c from *E. coli, Synechocystis* and *Sinapis* chloroplast shows structural similarity between cyanobacteria and chloroplasts, and dissimilarity between each and *E. coli* in residues upstream of the uS4c loop that borders the mRNA entrance channel (asterisk). **(C)** N-terminal extensions of uS5c contain unresolved loops (dashed lines) adjacent to the mRNA entrance channel that are in different locations in the chloroplast ribosomes of land plants (*Sinapis*) and green algae (*Chlamydomonas*). **(D)** Comparison of structural features on the ribosome surface bordering the mRNA entrance channel of the chloroplast and cyanobacterial ribosomes. Large domains on the surface of the *Chlamydomonas* chloroplast ribosome comprised of uS2c, uS3c, uS5c and PSRP7 (13) are absent from the chloroplast ribosome of *Sinapis* and other land plants. **(E)** Structural conservation of the ribosome decoding center. In all panels, structures are shown for *E. coli* (PDB:7K00) and *Chlamydomonas* chloroplast ribosome (PDB:9TVU) (13).

By contrast, the uS4c region upstream of this loop is structurally equivalent in the chloroplast and cyanobacterial ribosomes, indicating this feature that was identified as unlike *E. coli* (12) is not unique to the chloroplast (Figure 5B). The mRNA entrance channel of the chloroplast ribosomes of both plants and algae is also bordered by unresolved loops of uS5^NTE^ (Figure 5C). The mRNA entrance channel of land plant chloroplasts notably lacks the additional structural features comprising uS3c, uS4c, uS5c and cS25 observed in the *Chlamydomonas* ribosome (Figure 5D) (13).

### Conservation of the decoding and peptidyl transferase centers

The structure of the decoding centers of the *Synechocystis* and *Sinapis* chloroplast ribosomes closely resembles that of *E. coli* (Figure 5E). The P-site tRNA and mRNA are positioned for pairing by uS9/uS9c residues in common with *E. coli*, and 16S nucleotides with the same sequences and modifications. The peptide transferase center is similarly structurally conserved among *Sinapis, Synechocystis* and *E. coli*. We observed that the N-terminus of bL27/bL27c becomes ordered in translating states due to interactions with the A- and P-site tRNAs as in *E. coli*.

### Mechanisms of ribosome hibernation in chloroplasts and cyanobacteria

Non-translating ribosomes isolated from *Sinapis* chloroplasts and *Synechocystis* were associated with hibernation factors of the HPF family: chloroplast PSRP1 and cyanobacterial LrtA (Supplementary Figures S1H,S2G). This is consistent with evidence for the roles of HPF proteins in ribosome hibernation in cyanobacteria and the chloroplasts of plants and green algae (6, 10, 12, 13, 45-47). A subset of non-translating *Sinapis* ribosomes (15%) was additionally associated with RRF (Figure 6A, Supplementary Figure S1H), as previously characterized in non-translating spinach ribosomes (12). A subset of hibernating *Synechocystis* ribosomes (10%) were additionally associated with a homolog of the bacterial hibernation factor Balon (Figure 6A, Supplementary Figure S2G). The presence of Balon is consistent with the identification of Balon proteins in diverse bacteria following its recent discovery as a hibernation factor in *Psychrobacter* (48).

**Figure 6:**
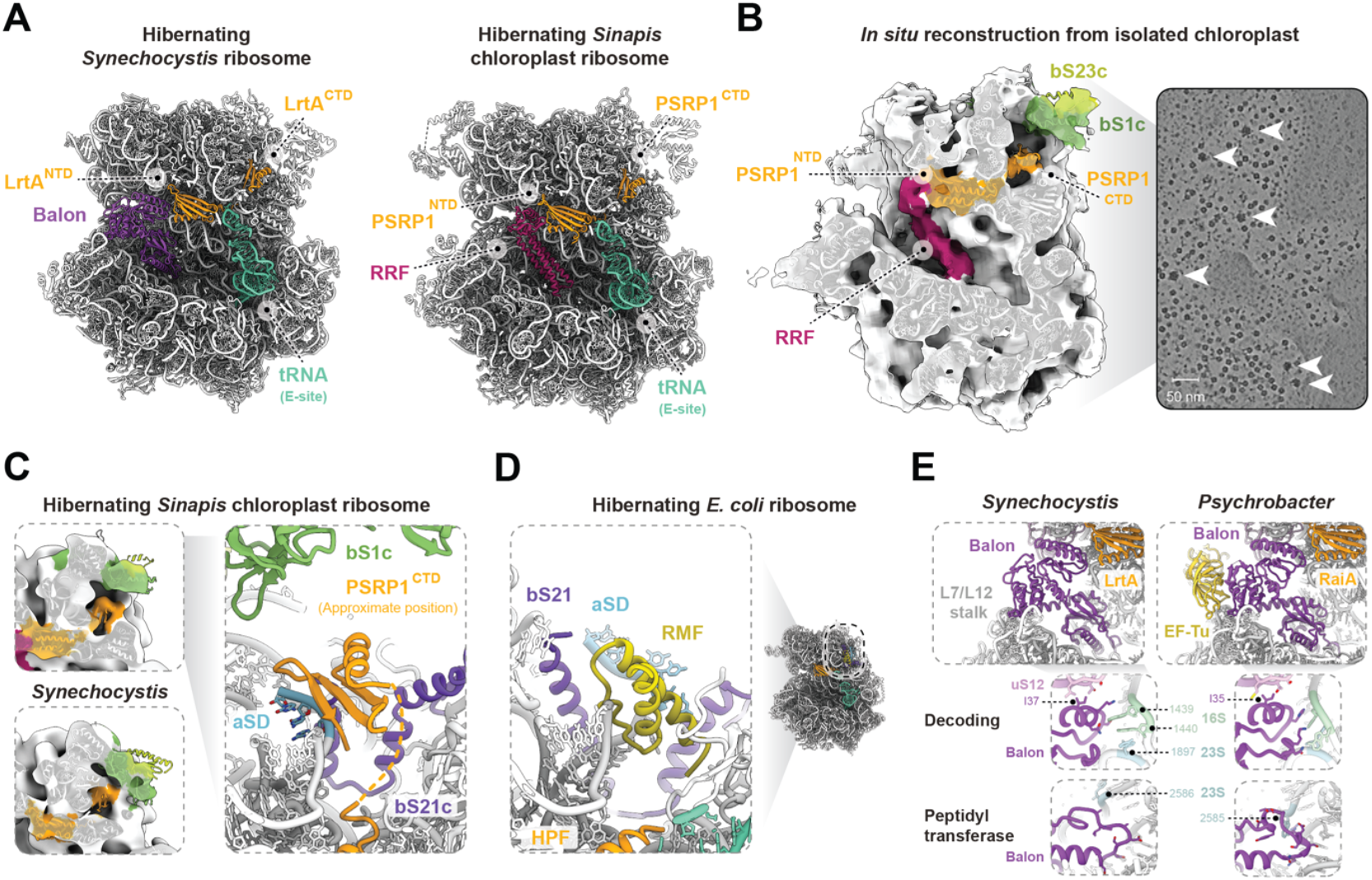
Mechanisms of ribosome hibernation in chloroplasts and cyanobacteria and schematic of evolutionary changes to ribosome structure in the photosynthetic lineage. **(A)** Structures of hibernating *Synechocystis* and *Sinapis* chloroplast ribosomes showing conserved positions of hibernation factors of the HPF family (orange), and similar binding sites of Balon and RRF in the cyanobacterial and chloroplast ribosomes respectively. **(B)** Left: Subtomogram average of spinach chloroplast ribosome bound in non-translating state bound to PSRP1 and RRF. Right: Slice of a tomogram with chloroplast ribosomes indicated with arrows. **(C)** Left: Cryo-EM reconstructions filtered to 16Å showing density in the 30S platform cavity consistent with HPF CTD domains. Right: The approximate position of the PSRP1 CTD showing the potential to prevent interactions between mRNA and the anti-Shine-Dalgarno (aSD) motif. **(D)** Structure of the *E. coli* ribosome bound to hibernation factors HPF and RMF (PDB:6H4N) (53), showing similar position of RMF to the PSRP1 CTD (panel C). **(E)** Structural similarity of Balon in hibernating ribosomes of *Synechocystis* and *Psychrobacter* (PDB:8RD8) (48).

To establish whether non-translating chloroplast ribosomes are bound by PSRP1, RRF or both prior to sample isolation we performed sub-tomogram averaging of ribosomes in a cryo-ET dataset of isolated spinach chloroplasts (49). A consensus in situ reconstruction was resolved to approximately 16 Å (Figure 6B, Supplementary Figure S7A-C). Classification revealed almost all analyzed ribosomes were not translating based on absence of tRNAs in the A- and P-sites (Supplementary Figure S7D). The cryo-EM density for RRF and PSRP1 indicates their occupancy is high and confirms both are associated in plant chloroplasts prior isolation. The state observed *in situ* differs from that of isolated *Sinapis* ribosomes in that E-site tRNA was not typically present (3% in chloroplasts compared to 87% isolated) (Figure S1H, Supplementary Figure S7D). Conversely, the occupancy of RRF was substantially higher *in situ* than following isolation (97% in situ compared to 15% isolated). This may reflect differences between species or leaf maturity, but it is also likely that RRF dissociation and tRNA association with the E-site occurred during isolation of ribosomes.

The cyanobacteria and chloroplast HPF proteins comprise two domains. In each, the N-terminal domain contacts the decoding center similarly to homologs in bacteria, *Chlamydomonas* chloroplasts, and spinach chloroplasts (Supplementary Figure S7E) (6, 10, 12, 45, 50, 51). We identified that the C-terminal domain (CTD) of PSRP1 and LrtA could be placed within poorly resolved density within the 30S platform cavity capped by bS1/bS1c in the cryo-ET reconstruction and filtered single-particle reconstructions (Figure 6B,C). The location and inconsistent orientation of the CTDs in each reconstruction suggest they are sterically trapped between the RNA-binding surfaces of bS21c and bS1c in a way expected to prevent Shine-Dalgarno duplex formation (Figure 6C).

HPF homologs in *E. coli* lack a CTD. Shine-Dalgarno duplex formation can instead be occluded during ribosome hibernation by ribosome modulation factor (RMF), a protein with homologs only in proteobacteria (50, 52, 53). We therefore speculate that the CTD-containing HPFs LrtA and PSRP1 perform the dual functions of *E. coli* HPF and RMF, occluding both the decoding center, like HPF, and the anti-Shine Dalgarno motif, like RMF (Figure 6D). *Staphylococcus aureus* HPF contains a CTD that mediates ribosome dimerization and is predicted to structurally resemble those of LrtA and PSRP1 (51). We did not observe ribosome dimerization in non-translating *Sinapis* or *Synechocystis* ribosomes despite prior evidence for hibernating ribosome dimerization in cyanobacteria (47) and the potential for dimerization observed in structural predictions (Supplementary Figure S7F).

Balon adopts a similar conformation in hibernating *Synechocystis* ribosomes to that in *Psychrobacter* despite sequence identity of only 30% (Figure 6E). Interactions with uS12 in the decoding center and the L7/L12 stalk are structurally similar, as predicted by the high relative conservation of Balon residues involved (48). By contrast, *Synechocystis* Balon makes extensive contact with rRNA in the decoding and peptidyl transferase centers through residues that are unlike the corresponding positions in *Psychrobacter* (Figure 6E). *Synechocystis* Balon does not contact LrtA, suggesting binding of each hibernation factor is independent, as in *Psychrobacter* (48). However, *Synechocystis* ribosomes containing Balon were consistently bound by LrtA, suggesting Balon selectively binds a 70S conformational state stabilized by LrtA. We did not observe the interactions identified in *Psychrobacter* between EF-Tu and Balon in *Synechocystis* (48), or between Balon and translating ribosomes.

## Discussion

Despite their common ancestry, chloroplasts differ from cyanobacteria in ways that could be expected to have driven evolutionary changes to the ribosome: a 10- to 100-fold reduced set of encoded proteins, and translation regulatory pathways linked to plant development and adaptation that are driven by proteins of eukaryotic origin imported to the chloroplast (2). Despite this, the ribosomes of chloroplasts and cyanobacteria are strikingly similar given the timespan since endosymbiosis. Through direct structural comparisons of the ribosomes of cyanobacteria and land plant chloroplasts, we show that the extent of similarity is even greater than previously thought. Features of the chloroplast ribosome of land plants inherited from its cyanobacterial ancestor include a homolog of bS23 (Figure 2A), the absence of bL25 (Figure S3A), bS1 and bS21 architecture associated with initiation (Figure 2A,D), uL24 features at the exit tunnel pore (Figure 3B), a largely conserved set of rRNA modifications (Figure S3), rRNA secondary structure elements (Figure 4C,G), and a structurally equivalent hibernation mechanism involving HPFs (Figure 6A).

By analyzing sequences of diverse plants, alga and bacteria, alongside structural comparison to the chloroplast ribosome of a green alga (13), changes potentially associated with endosymbiogenesis can be distinguished from those linked with the evolution of land plants (Figure 7). The conserved presence of cL38 and absence of bS24 among land plants and green algae indicate these changes may have occurred during endosymbiogenesis (Figure 4A). By contrast, cL37, cS22 and bTHXc (Figure 4C,E,G) arose in a common ancestor of land plants, along with a large set of proteins that supported terrestrialization (54). We observed that the terminal extensions of green alga chloroplast ribosomal proteins have little similarity to those of land plants and generally interact with distinct surfaces of the ribosome (Figure 2C, Supplementary Figure S4E). Thus, the presence of terminal extensions, but not their precise structural role, is a distinguishing feature of chloroplast ribosomes.

**Figure 7:**
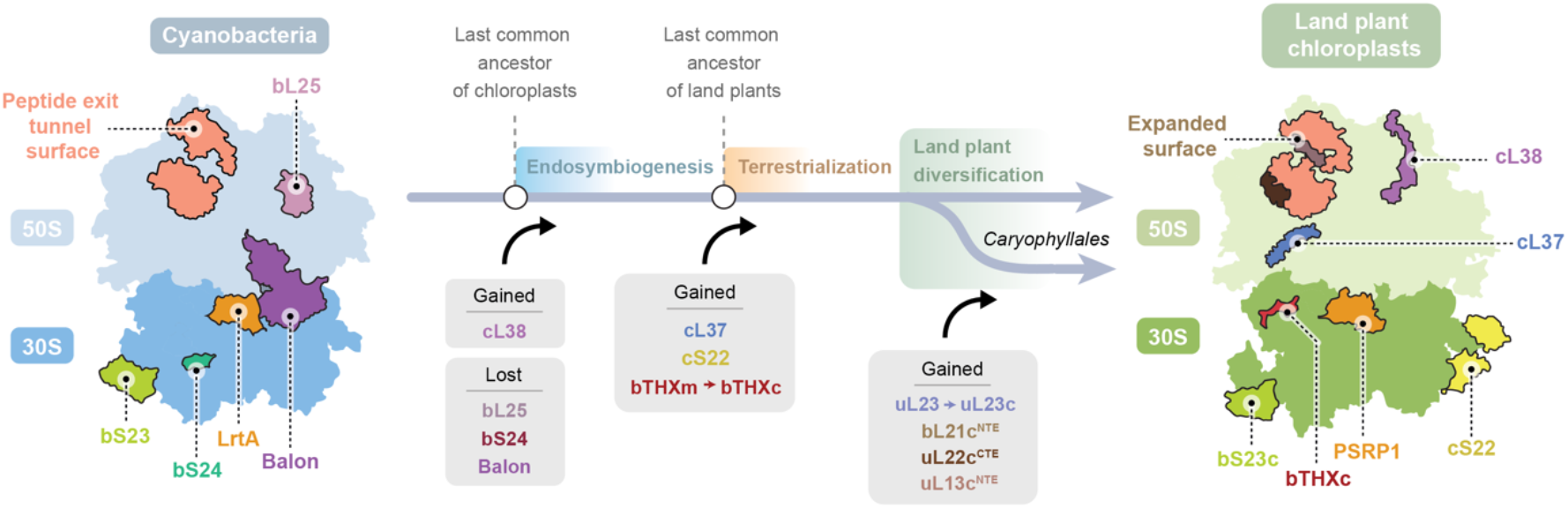
Evolutionary changes to ribosome composition in the photosynthetic lineage. Schematic representation of the gain and loss of ribosomal proteins mapped to the likely key evolutionary transitions between cyanobacteria and chloroplasts.

We identified that some ribosome features previously attributed as unique to the chloroplast are unusual features of the *Caryophyllales* clade of plants that includes spinach and are not shared by most other plants. In land plants other than *Caryophyllales*, the ribosomal proteins that surround the peptide exit tunnel (uL13c, bL21c, uL22c and uL23c) closely resemble cyanobacteria (Figure 3). The possibility that variation in chloroplast ribosome structures support distinct photosynthetic protein biogenesis pathways across plant species remains unexplored. The bS18c^NTE^, uL22c^CTE^ and uL29c^CTE^ display hallmarks of this potential (Figure 2F,3C). Terminal extensions in bS18c and uL22c are notable in that the presence of a terminal extension is uncommon in the plastid-encoded subset of ribosomal proteins. The increase in bS18c^NTE^ length from approximately 19 to 55 residues in the *Poales* clade of monocots may represent evidence of positive selection associated with a role in mRNA binding during translation initiation.

The evolutionary history of the chloroplast ribosome involves multiple examples of addition or replacement of ribosomal proteins. The incorporation of a uL23 homolog derived from the cytosolic ribosome across *Caryophyllales* accounts for the widespread pseudogenization of the plastid uL23c gene in this family (55). We further identify that bTHXc arose from the incorporation of a mitochondrial bTHX homolog in a common ancestor of land plants (Figure 4E). bTHX is a relatively uncommon component of bacterial ribosomes, and its ubiquitous presence only in *Thermales* and *Bacteroidetes* (18) suggests it may be advantageous to the tolerance of high temperatures. The role of bTHXc in plant organellar ribosomes is related based on structural similarity (Figure 4E,F). However, it must overcome an impediment to ribosome assembly or stability other than extreme temperature, such as the challenge to ribosome biogenesis of coordinating products of both the nucleus and organellar genomes. bTHXc is not unique among chloroplast ribosomal proteins in its non-endosymbiotic ancestry. bS16c and bL21c can similarly be encoded by nuclear genes of mitochondrial origin (56, 57). Conversely, mitochondrial ribosomal proteins uS13m and uL10m are evolutionarily derived from chloroplast genes. Swapping of ribosomal proteins among organelles has therefore occurred extensively through plant evolution (58, 59). Our data also confirmed the evolutionarily recent substitution of bS16c nuclear and chloroplast gene products within the brassica family (56).

We identified changes to the molecular basis of ribosome hibernation in the photosynthetic lineage. The association of HPF is conserved among cyanobacteria and the chloroplasts of land plants and green algae (Figure 6). By contrast, Balon, which can bind non-translating cyanobacterial ribosomes, appears to have been lost during endosymbiogenesis. In land plant chloroplasts, RRF contacts equivalent sites of the ribosome (Figure 6A), potentially suggesting a degree of mechanistic equivalence between Balon and RRF. To date, reactivation of ribosomes by eviction of HPF is a mechanism proposed for chloroplast RRF (6) but not Balon. However, Balon structurally resembles eukaryotic Pelota (48), which recruits a factor that rescues stalled ribosomes by nucleotide hydrolysis in a mechanism with parallels to the recruitment of EF-G by chloroplast RRF to release PSRP1 (6, 60). Thus, the basis of ribosome hibernation and reactivation by chloroplast RRF and PSRP1 could be informed by study of bacterial and eukaryotic processes. RRF was not observed bound to isolated *Chlamydomonas* chloroplast ribosomes (13), and it remains unclear if this role is conserved across the plant kingdom.

## Acknowledgements

We acknowledge Diamond for access and support of the cryo-EM facilities at the UK national electron Bio-Imaging Centre (eBIC). We thank Wojciech Wietrzynski for acquiring and curating spinach cryo-ET data. We thank JIC horticultural service provision for assistance with obtaining plant material. We thank all members of the Webster group for critical discussions.

## Author contributions

- Conceptualization: MWW, FW, DLS, BDE
- Methodology: DP, NNT, WTB, SD, IP, VKV, ATG, GG, ARJC, BM, JR, SR, GS, FW, MWW, BDE, DLS
- Investigation: DP, NNT, WTB, SD, IP, VKV, ATG, GG, ARJC, BM, JR, SR, GS, FW, MWW, BDE, DLS
- Visualization: DP, NNT, WTB, SD, MWW
- Funding acquisition: MWW, FW, DLS, BDE
- Project administration: MWW, FW, DLS, BDE
- Supervision: MWW, FW, DLS, BDE
- Writing – review & editing: NNT, WTB, ATG, MWW

## Data and code availability

Cryo-EM electron density maps and atomic coordinates have been deposited in the Electron Microscopy Data Bank (https://www.ebi.ac.uk/pdbe/emdb/) and RCSB Protein Data Bank (http://www.rcsb.org). PDB accession numbers for the *Synechocystis* ribosome are pdb_000031TT, pdb_000031TU, pdb_000031TV, pdb_000031TW and pdb_000031TX for the composite model, the translating state with three tRNAs, the translating state with EF-Tu, the non-translating state with EF-Tu and the non-translating state with Balon respectively. PDB accession numbers for the Sinapis chloroplast ribosome are pdb_000031VK, pdb_000031VL, pdb_000031VN for the composite model, the translating state and the non-translating state respectively. EMDB accession numbers are listed in Table S1. Mass spectrometry data have been deposited to the ProteomeXchange 403 Consortium via the PRIDE partner repository with the dataset identifier PXD080242.

## Lead contact

Correspondence should be addressed to

## Materials availability

Materials available upon request from Michael W. Webster.

## Materials and Methods

### Preparation of *Sinapis* chloroplasts

*Sinapis alba* was selected for analysis as it is fast growing and produces robust chloroplasts; features useful for isolation of large quantities of chloroplast protein complexes for structural analysis (1, 2). *Sinapis* is a member of the Brassicaceae family, which also includes the model species *Arabidopsis thaliana*, and is a superrosid, unlike the superasterid spinach, allowing structural conservation among eudicots to be assessed.

*Sinapis alba* (‘green manure’ from Moles Seeds) was sown in John Innes F2 Starter media and aerial parts of 1-week old seedlings were harvested. The subsequent steps were performed in a cold room at ∼8°C or in centrifuges operated at 4°C. Batches of 50 g *Sinapis* material were homogenized in 200 mL homogenization buffer comprised of 450 mM Sorbitol, 20 mM Tricine-KOH (pH 8.4 at 4°C), 10 mM EDTA, 10 mM MgCl_2_, 10 mM Na_2_CO_3_, 0.1% BSA, 1 mM DTT using a Waring blender in two pulses of 3 sec separated by 1 min. Homogenate was filtered through BioDesign cheesecloth and two layers of Miracloth. Chloroplasts were pelleted by centrifugation (1300×g, 10 min, 4 °C, rotor Fiberlite F9-6×1000 LEX) and material from each batch was resuspended in 5 mL resuspension buffer comprised of 300 mM Sorbitol, 20 mM Tricine-KOH (pH 7.6 at 4°C), 10 mM MgCl_2_, 2.5 mM EDTA, 1 mM DTT.

10 mL volumes of resuspended sample were loaded onto Percoll gradients, each prepared in a 30 mL Corex tube and comprised of 9 mL of 85% Percoll in resuspension buffer overlaid with 13 mL of 40% Percoll in resuspension buffer. Loaded Percoll gradients were centrifuged at 3214×g for 10 min at 4°C (Eppendorf 5810/5810R, S-4-104 rotor with breaking and acceleration settings of 0). Intact chloroplasts in the lower band were isolated with a 1 mL pipette with the tip removed following removal of the Percoll solution containing broken chloroplasts above. The intact chloroplasts were washed with 3 volumes of washing buffer comprised of 150 mM Sorbitol, 20 mM Tricine-KOH (pH 7.6), 10 mM MgCl_2_, 2.5 mM EDTA and centrifuged at 3214×g for 10 mins at 4°C (Eppendorf 5810/5810R, S-4-104 rotor). Approximately 5 mg of intact chloroplast was obtained per g of plant material, which was flash frozen in liquid nitrogen and stored at -80°C.

### Preparation of *Synechocystis* cells

*Synechocystis* sp. PCC 6803 cultures were grown in BG11 medium supplemented with 10 mM NaHCO_3_ at 30°C under continuous ∼50 µE m^-2^ s^-1^ light and with continuous shaking at 170 rpm. BG11 agar plates were prepared with BG11 medium supplemented with agar (15 g/L), TES (10 µM) and sodium thiosulfate (19 mM). *Synechocystis* sp. PCC 6803 starter cultures were prepared by inoculation of 50 mL BG11 from agar plates and were grown for 3-4 days. Growth cultures were prepared by inoculation of 1L of BG11 in 2L conical flasks with 50 mL of starter culture and incubated until the optical density at 750 nm (OD_750_) of the cultures was 0.7-0.9 (approximately 5-6 days). Cells were harvested by centrifugation at 7000×g for 20 mins at 22°C in a JLA 8.1000 rotor in a Beckman J-26 centrifuge. Pellets were resuspended in 10 mL BG11 using a Pasteur pipette and transferred to a pre-weighed 50 mL centrifuge tube. Cells were pelleted again by centrifugation at 3200×g for 20 minutes at 22°C. The remaining supernatant was removed with a pipette, with additional centrifugation where necessary. Tubes were weighed to determine cell mass, with a typical yield of ∼600 mg cells per 1 L culture, and stored at -80°C.

### Ribosome isolation by RAPPL

For the sample imaged in cryo-EM dataset 1, frozen intact *Sinapis* chloroplast pellet (3.6 g) was resuspended in non-detergent lysis buffer (23.5 mL) containing 20 mM HEPES-KOH (pH 7.5 at 4°C), 100 mM NH_4_Cl, 10.5 mM MgCl_2_, 0.5 mM EDTA, 1 mM CaCl_2_, 1 mM DTT, DNase I (4U/ml), Roche Complete protease inhibitor tablet (1 per 25 ml), RNaseOUT (40 U/mL), SUPERasIn (20U/mL). Chloroplasts were lysed by three passes through an Avestin EmulsiFlex-B15 homogenizer operated at 20,000 psi. Lysate was clarified by centrifugation at 3214×g for 10 mins at 4°C (Eppendorf 5810/5810R, S-4-104 rotor) and the supernatant filtered with a 0.45 µm PES syringe filter to remove debris. For the sample imaged in cryo-EM dataset 2, the *Sinapis* chloroplast pellet (3.4 g) was resuspended in the same lysis buffer (22 mL) with the addition of 2% Triton-X100. The sample was incubated for 40 min at 4°C with stirring before clarification and filtering rather than applied to a homogenizer. *Synechocystis* lysis was performed by passing cell pellet (2.3 g) resuspended in non-detergent lysis buffer (16 mL) through three passes of an Avestin EmulsiFlex-B15 homogenizer operated at 20,000 psi, before clarification and filtering.

Defrosted lysate (10 mL for non-detergent lysates or 8 mL for detergent lysate) was aliquoted in 1.25 mL volumes in Protein LoBind Tubes (Eppendorf) and clarified by centrifugation at 18000×g for 6 mins at 4°C in an Eppendorf centrifuge 5424R. 0.2 mL of polylysine magnetic beads (Molecular Cloning Laboratories) was added to each 1.25 mL aliquot of clarified lysate in new Protein LoBind Tubes. The samples were incubated with gentle agitation for 30 minutes at 4°C before supernatant was collected by pipette following magnetic immobilization of beads. Beads were washed three times with RAPPL wash buffer containing 20 mM HEPES-KOH (pH 7.5 at 4°C), 100 mM NH_4_Cl, 10.5 mM MgCl_2_, 0.5 mM EDTA and 1 mM DTT by repeated magnetic immobilization.

Ribosome samples were eluted from the beads for electron microscopy by incubation with 0.2 mL elution buffer comprising RAPPL wash buffer supplemented with poly-D-glutamic acid (Sigma; 2 mg/mL), 2.5 mM spermidine, Roche Complete protease inhibitor tablet (1 per 25 ml), RNaseOUT (40 U/mL) and SUPERasIn (20U/mL) for 15 mins at 4°C with gentle agitation before magnetic immobilization. The eluate was filtered with a 0.22 µm CA Spin-X Centrifuge Tube Filters (Costar) equilibrated with RAPPL wash buffer at 16000×g for 2 mins at 4°C to remove residual beads. Filtered eluate aliquots were pooled and concentrated using a VivaSpin 500 100,000 MWCO concentrator equilibrated with RAPPL wash buffer centrifuged at 12,000×g at 4°C to 0.25 mL for the *Sinapis* detergent lysate or 0.15 mL for *Synechocystis* and *Sinapis* non-detergent lysates. The buffer was exchanged to RAPPL wash buffer in the same concentrator. The sample analyzed in *Sinapis* cryo-EM dataset 1 was flash frozen in liquid nitrogen and defrosted for cryo-EM specimen preparation, whereas the sample analyzed in *Sinapis* cryo-EM dataset 2 and the *Synechocystis* sample were not frozen prior to cryo-EM specimen preparation.

For proteomic analysis of samples prepared by RAPPL, after the third wash of the magnetic beads, 0.1 mL of RAPPL wash buffer supplemented with 2% SDS was added to each tube. Samples were incubated at 90°C for 5 min to release proteins from the beads. Following centrifugation at 17000×g for 10 mins at 4°C in an Eppendorf centrifuge 5424R, the supernatant was collected and added to 0.6 mL of a mixture of methanol and chloroform (4:1 ratio). The solution was incubated on ice for 30 min and centrifuged at 17000×g for 15 mins at 4°C. The supernatant was carefully removed, and pelleted protein was washed with 1 mL of methanol. Following incubation at 4°C for 5 min, the sample was centrifuged at 17000×g for 10 mins at 4°C and the methanol removed. The sample was washed with another 1 mL of methanol and then washed once with 1 mL of acetone. The pellets were allowed to air-dry in a fume hood before processing of samples for mass spectrometry analysis.

### Negative stain electron microscopy

Negative stain electron microscopy was used to assess the composition of samples prepared by RAPPL. Grids coated with thin carbon (Electron Microscopy Sciences, #CF400-CU-50) were glow-discharged for 30 sec at 8 mA with a Leica ACE200 vacuum coater. 3.5 µL of diluted sample was applied to grids and incubated for 2 mins. Following blotting of excess solution, grids were stained with uranyl acetate solution (2% w/v) for 30 sec, before blotting again. 1059 micrographs were collected at room temperature on a Talos F200C transmission electron microscope (Thermo Fisher Scientific) operated at 200 keV using a Falcon 4i camera with settings: calibrated pixel size 2.0 Å/pixel, defocus −0.7 µm and 30 electrons/Å^2^. Data processing was performed with CryoSPARC. Particles were extracted with a box size of 288px binned to 72px for monosomes or 640px binned to 160px for polysomes for generation of two-dimensional class averages.

### Cryo-EM specimen preparation and data acquisition

UltrAuFoil R2/2 (Quantifoil, GmbH Germany) 200 mesh grids were plasma cleaned using a Harrick Plasma Cleaner for 45 seconds at high power. CHAPSO was added to a final concentration of 8 mM immediately before freezing for the sample from *Synechocystis* and one of the two samples from *Sinapis* chloroplasts. 3.5µl of sample prepared by RAPPL was applied to the grid in a Vitrobot Mark IV plunger (FEI) at 4ºC and 100% humidity and vitrified by plunging into liquid ethane. Cryo-EM micrographs were collected using a Titan Krios transmission electron microscope (FEI) operated at 300 keV with a BioQuantum energy filter (slit width 20 eV) and K3 summit direct electron detector (Gatan).

### Cryo-EM Data Processing

Cryo-EM data was processed with CryoSPARC version 4 and reconstructions were interpreted with ChimeraX version 1.10. Each dataset was initially processed with the ‘Patch Motion Correction’, ‘Patch CTF’ and ‘Exposure Curation’ jobs. This resulting number of micrographs used is indicated in Table S1. Particles were picked with the ‘Template Picker’ job using 29 class averages from a preliminary dataset and settings of 20Å template filter, 20Å angular sampling of templates, and local maxima limit of 500. Particles were extracted (Figures S1,S2) with 2× binning (512px reduced to 256px), and correct picks were selected using a ‘Heterogeneous Refinement’ job with an initial reconstruction of the chloroplast ribosome and six ‘decoy’ reconstructions generated using an ‘Ab-Initio’ job. A further round of ‘Ab-initio’ and ‘Heterogeneous Refinement’ jobs were used to select particles assigned to a well-resolved reconstruction. These particles were re-extracted without binning (640px) and processed by a ‘Reference Motion Correction’ job following initial ‘Homogeneous Refinement’. The consensus 70S reconstruction was generated using a ‘Homogeneous Refinement’ job with optimization of per-particle defocus and per-group tilt and trefoil CTF parameters and a mask around the entire 70S.

Resolvability was improved using a set of ‘Local Refinement’ jobs with focused masks around different regions of the 70S ribosome consensus reconstruction, with ‘Marginalization’ and ‘Non-uniform Refine’ options enabled. Masks were created using the ‘Molmap’ command in ChimeraX with preliminary atomic models with 10Å lowpass filtering applied, followed by mask generation with 15Å lowpass filtering, threshold 0.1, dilation of 4px and soft padding of 6px. First, local refinements were performed with masks around either the 50S or 30S ribosome subunits. Further focused refinement of the 30S subunit was performed with masks around either the 30S head, 30S shoulder or the 30S body and tail regions. Further focused refinement of the 50S was performed with 5 partially tiled overlapping region. Finally, reconstructions with better resolution of dynamic regions on the ribosome surface (bS1c and bS23c, cS22, uL1, uL10 and uL11) were obtained using focused classification jobs, in which particle subsets with more consistent positions of these regions were selected.

The 70S particle set aligned to the 50S were classified into translating and non-translating states using a focused mask around the A-, P- and E-tRNA sites and the uL1 stalk filtered to 15Å in a ‘3D Classification’ job. For the *Sinapis* chloroplast dataset prepared without CHAPSO, 1,116,911 particles were classified into the following states: non-translating 70S bound to PSRP1 and E-site tRNA (50% of particles; 627,439 particles), translating 70S with tRNAs in the A-, P- and E-site (25% of particles; 318,285 particles), and non-translating ribosomes bound to RRF, PSRP1 and E-site tRNA (10% of particles; 129,252 particles) (Figure S1H and Table S1). The remaining particles contained states of lower prevalence that were not modelled as they contain structural information that is redundant with the modelled states: non-translating 70S bound to PSRP1 but not E-site tRNA, and translating 70S without tRNAs bound in all three sites.

For the *Synechocystis* dataset, 510,191 particles were classified into the following states: non-translating 70S bound to LrtA, E-site tRNA and EF-Tu represented (52% of particles; 264,666 particles), non-translating 70S bound to LrtA, E-site tRNA and Balon represented (7% of particles; 33,746 particles), translating 70S with tRNAs in the A-, P- and E-site (7% of particles; 33,703 particles), translating 70S with P- and E-site tRNAs and EF-Tu (5% of particles; 24,875 particles) (Figure S2G, Table S1). The remaining particles either contained partially resolved density for tRNAs in translating 70S or were assigned to a poorly resolved 70S. For each state, reconstructions were obtained by ‘Local Refinement’ jobs with mask around either the 50S subunit or the 30S applied during refinement.

FSC information was estimated for each reconstruction using CryoSPARC. The local resolution of reconstructions was estimated using the job ‘Local Resolution’ and visualized in ChimeraX, for example using the command ‘color sample #1.1 map #2 palette 1.4, #e68fef:1.6, #b36fc9:1.8, #6e3ba7:2, #4b78e3:2.2, #79c01a:2.4, #fbea38:2.6, #ffaa03:2.8, #e81815:3, #ab0608:3.2, #82080e:4, #5b060b key true’ for the resolution range 1.4 to 4 (for map #1 and local resolution file #2), or the command ‘color sample #1.1 map #2 palette 2, #4b78e3:2.5, #79c01a:3, #fbea38:3.5, #ffaa03:4, #e81815:4.5, #ab0608:5, #82080e:6, #5b060b key true’ for the resolution range 2 to 6.

Composite cryo-EM reconstructions were generated from focused refinement reconstructions using ChimeraX. Focused reconstructions were aligned to the 70S consensus, for example using the ChimeraX command ‘fit #2 in #1’ repeated for #3 to #9 (for consensus #1 and focused reconstructions #2-9). Density outside the masked region used for local refinement was erased for each focused reconstruction using the ChimeraX ‘volume eraser’ tool. The composite map was generated using the ChimeraX ‘volume maximum’ command with resampling from ∼0.8 Å/px to 0.34 Å/px to suppress resolution loss arising from resampling, for example using the ChimeraX command ‘volume maximum #1-9 onGrid #1 spacing 0.38’ (for consensus #1 and focused reconstructions #2-9). The composite map was recentered in a smaller box to reduce file size, for example using the ChimeraX command ‘volume boxes #10 centers #11 size 280’ (for a marker #11 in the center of composite map #10). Finally, the origin was reset and the file saved, for example using the ChimeraX command ‘volume #12 origin 0,0,0; save “<name>” models #12’ (for composite map #12).

### Structural model building

An initial structural model of the *Sinapis* chloroplast ribosome was constructed by combining all rRNA chains from an *E. coli* ribosome structure determined from a reconstruction at 1.55Å (PDB:8B0X) (3) with protein chains from a spinach chloroplast ribosome structure determined from a reconstruction at 2.9-3.5Å (PDB:6ERI) (4). Protein residues were mutated to the sequences of the *Sinapis alba* protein isoform identified by LC-MS. An initial model of the *Synechocystis* ribosome was constructed from the model of the *Sinapis* chloroplast ribosome. Models for each protein chain was obtained from the AlphaFold database (5) and aligned to the *Sinapis* model, and models for each rRNA chain were generated by mutating the sequences of those in the *Sinapis* model. Cryo-EM density was assigned to Balon using automated model building of the map region with ModelAngelo (6), and an initial model of Balon was generated by aligning the model from the AlphaFold database to the ModelAngelo model. Cryo-EM density for tRNAs reflects the mixture of tRNA species that co-purified with the ribosome. The models include tRNA-Phe to represent the ensemble present at each tRNA position. Initial mRNA models were obtained from a structure of the translating *E. coli* ribosome (PDB:6ZTJ) (7). RNA sequences for *Sinapis* were obtained from genome sequence (accession: NC_045948.1) with the following nucleotide regions: 16S (100308-101798), 23S (103993-106802), 4.5S (130366-130264), 5S (107251-107371) and tRNA-Phe(GAA) (47126-47198). RNA sequences for *Synechocystis* were obtained from the CyanoCyc server (8): 16S (SGL-RS13310), 23S (SGL-RS13300), 5S (SGL-RS13295) and tRNA-Phe (SGL-RS09435). Refinement was performed with Coot-1.1.18 and Phenix version 2.0.5936.

### Cryo-ET data acquisition

CryoET data analysis was performed on a deposited cryo-ET dataset (EMPIAR-12612) (9). In brief, tilt-series of isolated *Spinacia oleracea* chloroplasts were collected with a 300kV Titan Krios TEM (Thermo Fisher Scientific), with a post-column energy filter (Quantum, Gatan), and a direct detector camera (K2 Summit, Gatan). SerialEM was used to acquire a dose-symmetric tilt-scheme (10) with 60 tilts with 2° increments. An imaging rate of 8-12 frames/second at a pixel size of 3.52 Å, and a defocus range of 2.5-5 µm were applied, resulting in a nominal dose of 2.0 e^-^/Å^2^ per tilt image.

### Cryo-ET subtomogram averaging

25 tomograms were selected from Wietrzynski *et al*. (9) and processed following methods described in the TomoGuide (11). AreTomo3 (12) was used to perform motion-correction, CTF-estimation and tilt-series alignment, and reconstruction of CTF-corrected tomograms (at 14.08 Å/pixel, check please) in an automated manner. Denoising was performed using IceCream (13) on tomograms reconstructed from odd and even frames.

Particle picking was performed using pytom-match-pick (14) on bin4, CTF-corrected tomograms using the single particle structure of the Spinach ribosome (EMD-6709) (15) as a template, with 10 degrees angular sampling. The number of extracted positions was automatically determined for each tomogram individually using the pytom built-in cut-off estimation, and 1670 particles were extracted in total. Subtomogram averaging was performed using RELION5.0. Particles were initially subjected to 3D classification, after which the best classes, representing 1511 particles were selected. We then iteratively performed alignment, per-particle CTF-estimation and Bayesian polishing of the tomograms to reach a resolution of 18.4 Å. Tomograms were then re-aligned based on refined particle positions using polish2aretomo3 (16), and template matching was performed in the same manner with the resulting bin4 average as a template.

At this point, 2513 particles were extracted as bin4 2D-stacks, and subjected to 3D classification without alignment to remove false positives, after which 2404 particles were retained. These particles were then aligned in bin4, and reached bin4 Nyquist (28.2 Å). From there, a bin1 reference was reconstructed for per-particle CTF-refinement and Bayesian polishing, which improved the overall resolution to 18.8 Å. 2D stacks were extracted at bin2, and subjected to another round of 3D refinement. 3D classification without alignment was used again to remove the last remaining false positives, resulting in a finalized particle list of 2377 particles. Another round of CTF-refinement, Bayesian polishing and alignment at bin1 finally resulted in an average of 16.1 Å overall resolution. Classification of the aligned particles in bin4, with a mask focused on the tRNA sites, yielded only two distinguishable classes, one fully without tRNA (2303 particles, 96.9%) and one with a density at the exit site (74 particles, 3.1%). These classes were reconstructed and post-processed in bin2 for comparison and visualization.

### Analysis of protein conservation

Protein conservation was assessed by sampling representatives from diverse *viridiplantae* taxa. Genomes were downloaded from NCBI (17), Phytozome (18), Phycocosm (19) and treegenesdb (20) alongside predicted proteome annotations when available. Genomes without annotations were annotated with Helixerv0.3.6 (21) with the option ‘--lineage land_plant’ applied to the default model. Annotations were also included from Toghani *et al*. (22), which were previously annotated by the same method. Genome annotations were assessed using BUSCO scores v5.50 (23) with options ‘-m protein --lineage_dataset viridiplantae_odb10’. The predicted proteome database comprised 809 species of land plants, algae, cyanobacteria, and bacteria.

Ribosomal protein homologs were identified in the database using HMMERv3.4 (24). Hits were extracted using custom R scripts and aligned using MAFFTv7.525 (25). Alignments of the full-length sequences and the domains were used to construct phylogenetic trees using VeryFastTree v4 (default settings) (26). The initial phylogeny was refined by extracting selected sequences from well-supported clades using CD-HIT v4.8.1 (27) with option ‘-c 1.0’ to remove redundant sequences. Sequences were aligned using MAFFTv7.525 (25) with options ‘--reorder --maxiterate 1000 –localpair --op 3 --ep 0.2’. Phylogenetic trees were drawn using FastTreev2.2 (28), rooted using the bacterial homolog clade, and visualized using iTOL v7.6 (29).

Conservation of cyanobacterial proteins was further assessed by targeted BLAST search for representatives within each clade of cyanobacteria defined by Moore *et al*. (30). Structural predictions were generated using the AlphaFold 3 webserver (31). AlphaFold models, protein sequence alignments and PAE plots were then imported into ChimeraX (32-34).

### Liquid chromatography mass spectrometry

Protein sample (estimated 20 µg) was added to 50 µL of 1.0% sodium deoxycholate (SDC; Merck) in 0.2 M EPPS-buffer (Merck), pH 8.5 and vortexed under heating. Samples were treated with TCEP and CAA to reduce cysteine residues and the proteins were digested with trypsin in the SDC buffer according to standard procedures. After the digest, the SDC was precipitated by adjusting to 0.5% trifluoroacetic acid (TFA), and the clear supernatant subjected to C18 SPE using stage tips (Affinisep, France). Aliquots were analyzed by nanoLC-MS/MS on an Orbitrap Eclipse Tribrid mass spectrometer with a Thermo Scientific FAIMS Pro Duo source and coupled to a Vanquish Neo UHPLC System (Thermo Fisher Scientific, Hemel Hempstead, UK). The samples were loaded onto a trap cartridge (PepMap− Neo Trap Cartridge, C18, 5um, 0.3×5mm, Thermo Fisher Scientific) with 0.1% TFA. The trap column was then switched in-line with the analytical column (Aurora Frontier TS, 60 cm nanoflow UHPLC column, ID 75 µm, reversed-phase C18, 1.7 µm, 120 Å; IonOpticks, Fitzroy, Australia) for separation at 60°C using the following gradient of solvents A (water, 0.1% formic acid) and B (80% acetonitrile, 0.1% formic acid) at a flow rate of 0.25 µL/min : 0-4 min 1%-6% B (curve 4); 4-154 min linear increase B 6%-154%, followed by a ramp to 99% B and re-equilibration to 1% B. Mass spectrometry data were acquired with the FAIMS device set to three compensation voltages (-35V, -50V, -65V) for 1 s each with temperatures of 80°C and 100°C for outer and inner electrodes in positive ion mode with the following MS settings: resolution 120K, profile mode, mass range m/z 300-1600, AGC target 4e5, max. inject time 50 ms; MS2: quadrupole isolation window 1 Da, charge states 2-5, threshold 1e4, HCD CE = 30, AGC target standard, max. injection time dynamic, dynamic exclusion 1 count for 15 s with mass tolerance of ±10 ppm.

The mass spectrometry raw data were processed and quantified in Proteome Discoverer 3.3 (Thermo Fisher Scientific) using the search engine CHIMERYS 5.8 (MSAID, Munich, Germany); all mentioned tools of the following workflow are nodes of the proprietary Proteome Discoverer (PD) software. Corresponding protein sequence databases (*Synechocystis*, Uniprot UP000001425, April 2026, 3507 entries; *Sinapis alba*, Salba_584_v3.1.protein, 54925 entries, including changes to PEP subunit sequences indicated in Table S6) was imported into PD adding a reversed sequence database for decoy searches; a database for common contaminants (maxquant.org, 245 entries) was also included. The database search was performed using the incorporated CHIMERYS (MSAID, Munich, Germany) with the inferys_4.7_fragmentation prediction model, a fragment tolerance of 0.3 Da, enzyme trypsin with 2 missed cleavages, variable modification oxidation (M), fixed modification carbamidomethyl (C) and with FDR targets 0.01 (strict) and 0.05 (relaxed). The workflow included the Minora Feature Detector with min. trace length 5, S/N 2.5, PSM confidence high; the Top N Peak Filter with 20 peaks per 100 Da. The consensus workflow in the PD software was used to evaluate the peptide identifications and to measure the abundances of the peptides based on the LC-peak intensities. For identification, an FDR of 0.01 was used as strict threshold, and 0.05 as relaxed threshold. For protein abundances the average of the top3 most abundant unique peptides was used. The results were exported into a Microsoft Excel table including data for protein abundances, number of peptides, protein coverage, q-values, and the CHIMERYS identification score.

---

**Figure S1:**
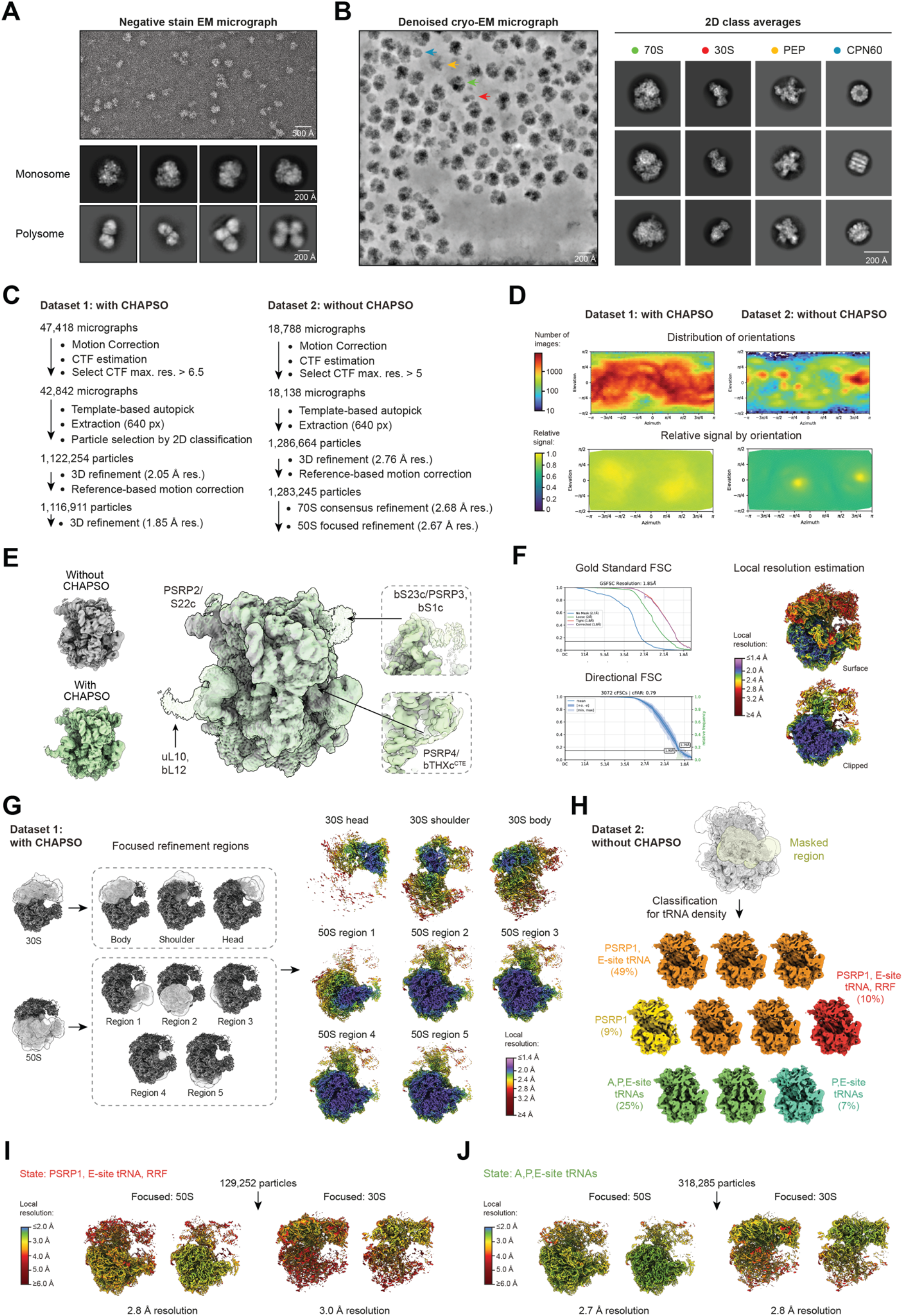
Cryo-EM analysis of Sinapis chloroplast ribosomes. **(A)** Analysis of RAPPL eluate from *Sinapis* lysate by negative stain EM. Representative micrograph and selected two-dimensional class averages of individual ribosomes (‘Monosome’) and ribosomes isolated translating the same endogenous mRNA (‘Polysome’) are shown. **(B)** Analysis of RAPPL eluate from *Sinapis* lysate by cryo-EM. Representative denoised micrograph (left) from the dataset collected in the absence of CHAPSO, with arrows indicating examples of the four constituents identified from two-dimensional class averages (right): 70S ribosome (green), 30S ribosomal subunit (red), plastid-encoded RNA polymerase (PEP; yellow), and CPN60 (blue). **(C)** Data processing workflows for two cryo-EM datasets used to characterize the *Sinapis* chloroplast ribosome. The dataset collected in the presence of CHAPSO (left) produced reconstructions of the highest resolution that were used to generate the composite map used for consensus model building, whereas the dataset collected in the absence of CHAPSO (right) produced reconstructions of distinct ribosome states. **(D)** Plots of angular distribution and relative signal by orientation indicate improved sphericity of the cryo-EM reconstructions obtained from the data collected in the presence of CHAPSO. **(E)** Comparison of low-pass filtered consensus cryo-EM reconstructions of the *Sinapis* chloroplast ribosome obtained from data collected in the presence (green) or absence (gray) of CHAPSO. Examples of cryo-EM density at the ribosome peripheries that are present only in the reconstructions obtained from data collected with CHAPSO include chloroplast-specific features. The protective effect of CHAPSO in cryo-EM sample preparation has been described (18, 19). **(F)** Fourier shell correlation (FSC) plots and local resolution estimates for consensus reconstruction of the *Sinapis* chloroplast ribosome. **(G)** Regions masked during focused refinement and classification (left) and resulting reconstructions colored by local resolution (right) combined to produce the composite reconstruction of the *Sinapis* chloroplast ribosome used for model building. **(H)** Scheme for focused classification of particle images with a mask around tRNA binding sites used to generate reconstructions of ribosome states. **(I**,**J)** Local resolution of reconstructions of non-translating (I) and translating (J) states of the *Sinapis* chloroplast ribosome focused on the 50S subunit (left) or 30S subunit (right).

**Figure S2:**
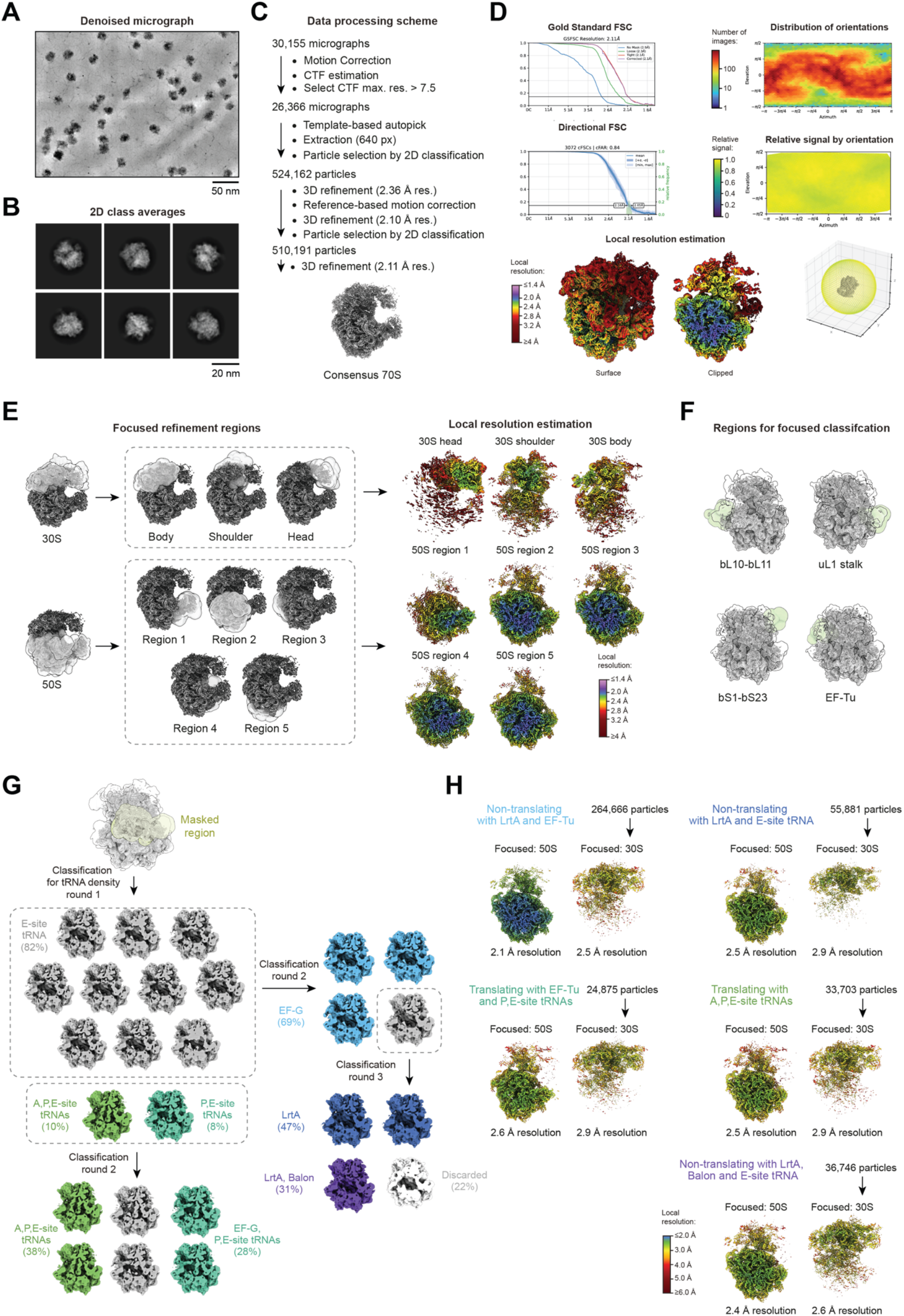
Cryo-EM analysis of *Synechocystis* sp. PCC 6803 ribosomes. **(A)** Representative denoised cryo-EM micrograph of *Synechocystis* ribosomes isolated by RAPPL. **(B)** Representative two-dimensional class averages. **(C)** Cryo-EM data processing workflow leading to consensus reconstruction of the *Synechocystis* ribosome. **(D)** Fourier Shell Correlation (FSC), angular distribution and relative signal by orientation plots and local resolution estimates for consensus reconstruction of the *Synechocystis* ribosome. **(E)** Regions masked during focused refinement and classification (left) and resulting reconstructions colored by local resolution (right) combined to produce the composite reconstruction of the *Synechocystis* ribosome used for model building. **(F)** Regions masked during focused classification to obtain reconstructions with improved resolution for dynamic surface regions included in the composite reconstruction of the *Synechocystis* ribosome. **(G)** Scheme for focused classification of particle images with a mask around tRNA binding sites used to generate reconstructions of ribosome states. **(H)** Local resolution of reconstructions of non-translating and translating states of the *Synechocystis* ribosome focused on the 50S subunit (left) or 30S subunit (right).

**Figure S3:**
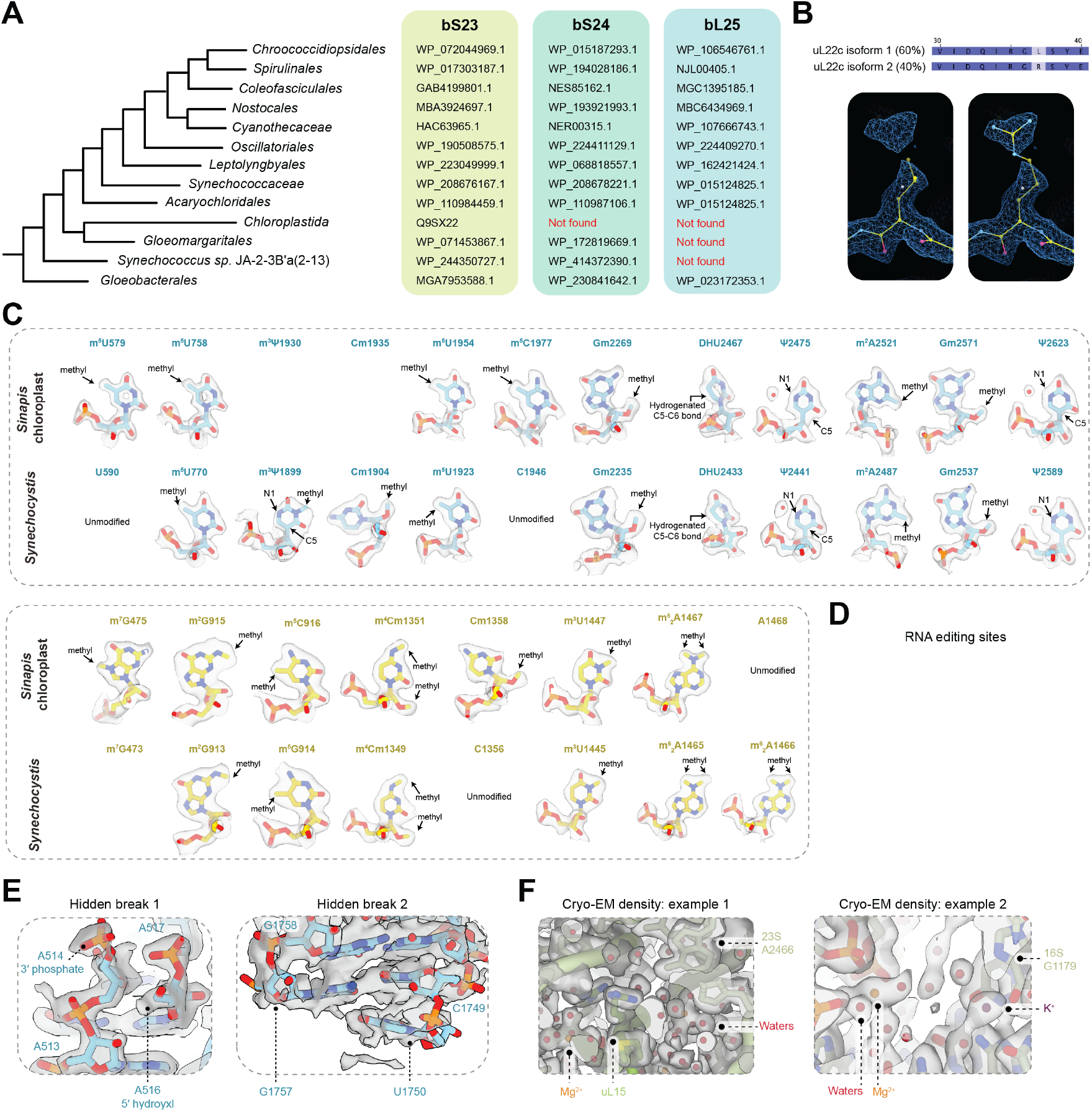
Composition and structural details of the cyanobacterial and chloroplast ribosomes. **(A)** Analysis of ribosomal protein composition across cyanobacteria and plant chloroplasts. Homologs of bS23, bS24 and bS25 were identified by reciprocal BLAST search using representative clades selected for maximum diversity (30). **(B)** Example of ribosome heterogeneity arising from presence of alternative isoforms. Top: relative of abundance of uL22c isoforms quantified by LC-MS. Bottom: cryo-EM density showing evidence for the overlapping presence of leucine and arginine due to isoform variation. **(C)** rRNA modifications of the cyanobacterial and chloroplast ribosomes in 23S (blue) and 16S (yellow). Model and cryo-EM density are shown for nucleotides at equivalent positions in the *Sinapis* chloroplast (top) and *Synechocystis* (bottom). Cytosine methylations Cm1356 (*Synechocystis*) and Cm1358 (*Sinapis*) occur at positions unmodified in *E. coli* but modified in *Mycobacterium tuberculosis* and *Arabidopsis* (37, 38). Cytosine methylations Cm1904 (*Synechocystis*) and Cm1920 (*Sinapis*) occur at positions unmodified in *E. coli* but modified in diverse bacteria, where it is catalyzed by homologs of TylA (37-41). Guanine methylations Gm2537 (*Synechocystis*) and Gm2571 (*Sinapis*) occur at positions unmodified in *E. coli* but modified in eukaryotic cytosolic ribosomes (38). **(D)** Cryo-EM density supports the presence of leucine in uS14c (residues 27 and 50) and uL23c (residue 30) at positions encoded as prolines and serines and altered by RNA editing (42-44). **(E)** Left: Cryo-EM density supports the presence of a hidden break in the 23S rRNA between nucleotides A514 and A516 as previously reported (45, 46). The improved resolution indicates the product of rRNA processing at this site produces a 3′ phosphate on the upstream strand. Right: Cryo-EM density supports the position of the second 23S rRNA hidden break between nucleotides A1756 and G1757 through the improved resolvability of G1757 relative to nucleotide A1756. **(F)** Examples of density in the *Sinapis* chloroplast ribosome reconstruction that supports the modelling of ions and waters.

**Figure S4:**
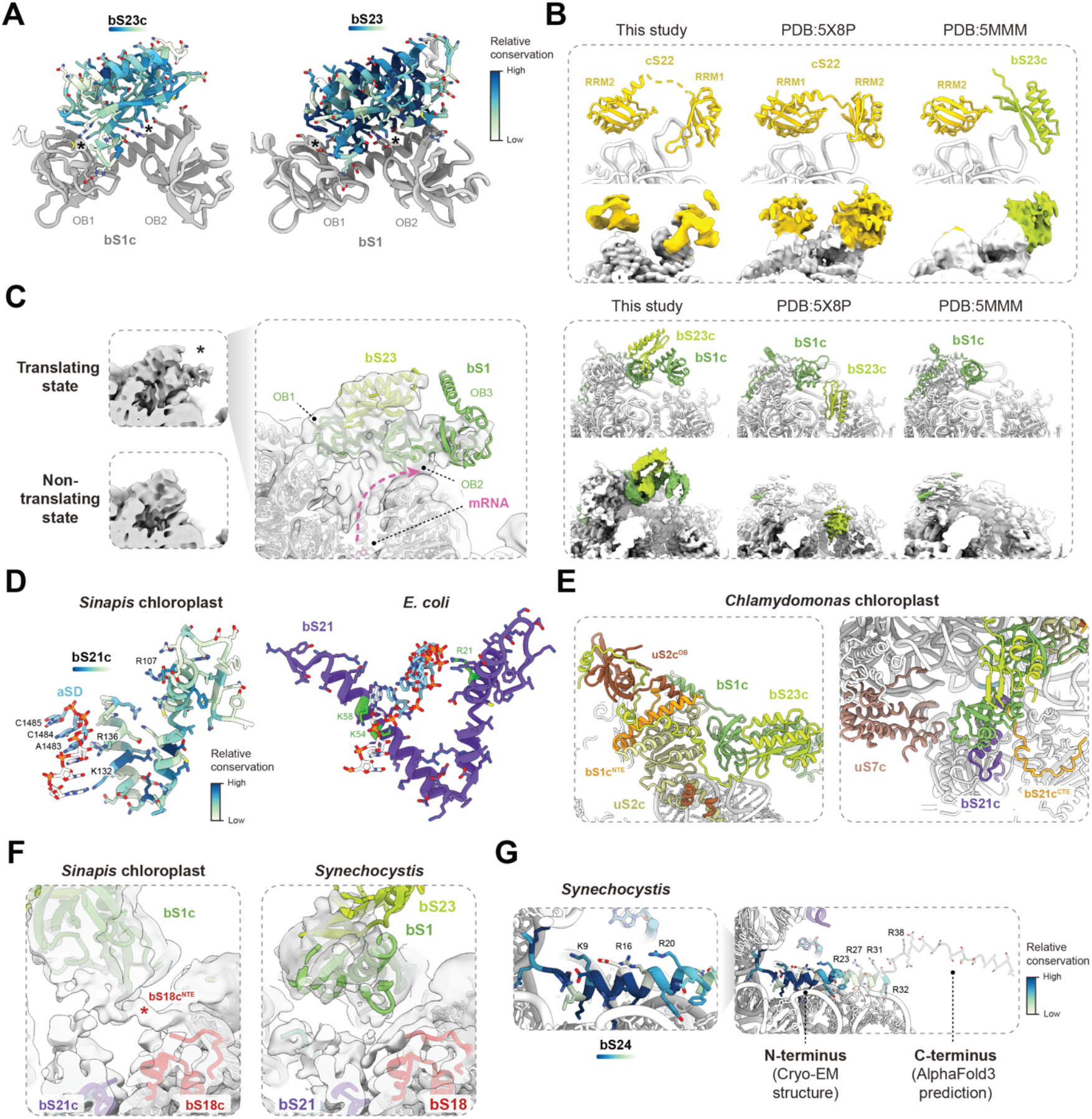
Additional data on the distinct ribosome features associated with translation initiation in the green lineage. **(A)** Residue-level conservation of bS23 and bS23c mapped onto structures of *Synechocystis* (left) or *Sinapis* chloroplast (right) shows conservation of residues at the interface with the bS1/bS1c OB1-OB2 linker (asterisks). **(B)** Assignment of bS23c/PSRP3 and cS22/PSRP2 in cryo-EM reconstructions. Comparison of models from the current study (left), PDB:5X8P (15) (middle) and PDB:5MMM (20) (right) with corresponding maps colored by protein assignment. Each map was filtered to 6Å for direct comparison (EMDB:6709 and EMDB:3533). Although the approximate position of PSRP2/cS22 at the foot of the 30S was correctly identified in models of the spinach chloroplast ribosome, the relative positions of its two RRM domains was not. The position of PSRP3/bS23c was incorrectly assigned to the 30S foot or 30S head in models of the spinach ribosome. **(C)** Comparison of reconstructions of translating and non-translating states of the *Synechocystis* ribosome (each filtered to 8Å) showing the approximate position of the bS1 OB3 domain. The consensus path of the exiting mRNA visible as poorly resolved density is indicated. **(D)** Residue-level conservation of bS21c mapped onto structure shows conservation of basic residues that contact the anti-Shine-Dalgarno (aSD). Comparison to *E. coli* translation initiation complex containing incoming mRNA (PDB:9GUT) (19) with basic residues contacting the aSD highlighted green. **(E)** The bS1c N-terminal extension (bS1c^NTE^) and bS21c C-terminal extension (bS21c^CTE^) are structurally different in the *Chlamydomonas* and *Sinapis* chloroplast ribosomes. Structure of the *Chlamydomonas* chloroplast ribosome (PDB:9TVU) (21) is shown in the views equivalent to Figures 2D and 2E. The OB domain present in algae chloroplast uS2c but not plant chloroplast uS2c (uS2c^OB^) is located distal to bS1c and PSRP3. **(F)** Cryo-EM reconstruction of the *Sinapis* chloroplast ribosome filtered to 4Å resolution shows density (asterisk) consistent with the dynamic N-terminal extension of bS18c (bS18c^NTE^). The absence of corresponding density in the reconstruction of *Synechocystis* filtered to 4Å resolution is consistent with the assignment to a chloroplast-specific sequence feature. **(G)** Residue-level conservation of bS24 mapped onto structure shows conservation of basic residues facing the 30S platform cavity. Structural prediction of *Synechocystis* bS24 was generated to visualize conservation of unmodelled residues inferred to by dynamic. The presence of additional basic residues of low conservation within the unstructured C-terminus is indicated.

**Figure S5:**
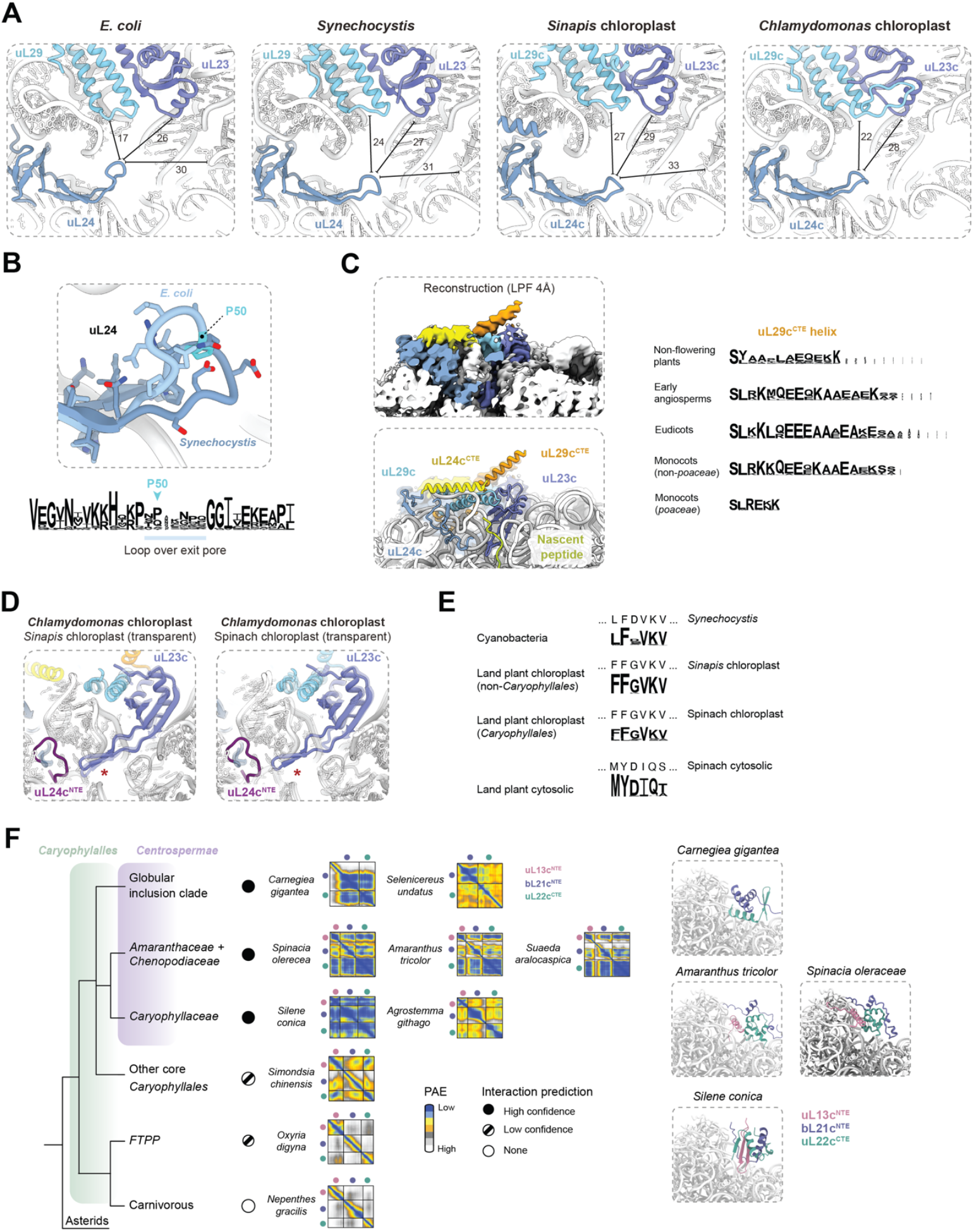
Additional data on the peptide exit channel in the green lineage. **(A)** Variation in peptide exit pore size across bacteria and chloroplasts. The pores of cyanobacteria and chloroplasts are larger than *E. coli* primarily due to the altered position of the uL24 loop overhanging the pore. The pore of the plant chloroplast is further largened relative to cyanobacteria due to a shift of uL29c by ∼3Å from the exit channel. Distances between Cα or O4′ atoms are shown in Å. **(B)** *E. coli* uL24 proline residue 50 produces a bend in the loop overhanging the peptide exit tunnel. Below: Conservation of uL24 across diverse bacteria shows variation in the loop that overhangs the exit tunnel pore. **(C)** Cryo-EM reconstruction of *Sinapis* chloroplast ribosome low-pass filtered to 4Å shows the positions of the C-terminal extensions of uL24 (uL24^CTE^) and uL29 (uL29^CTE^) that overhang the peptide exit pore. The sequence of the uL29^CTE^ C-terminal helix varies across plant clades. **(D)** uL23c in the *Chlamydomonas* peptide exit tunnel (opaque) is structurally similar to *Sinapis* (left, transparent) and distinct from spinach (right, transparent). The N-terminal extension of *Chlamydomonas* uL24 (uL24^CTE^) is absent from the peptide exit tunnel of land plant chloroplasts. **(E)** A loop of uL23c is conserved in sequence across chloroplast-targeted gene products despite the eukaryotic-origin of *Caryophyllales* uL23c. **(F)** Structural predictions of interactions between uL13c^NTE^, bL21c^NTE^ and uL22c^CTE^ across *Caryophyllales* clades. Plots of predicted alignment error (PAE) show interactions between sequence extensions for representatives from each clade, except for carnivorous *Caryophyllales*. Structural predictions for representative species were generated with AlphaFold3 and are overlayed in approximate locations to a structure of the spinach chloroplast ribosome (transparent, PDB:6ERI) (4), revealing diversity in the nature of the interactions between sequence extensions.

**Figure S6:**
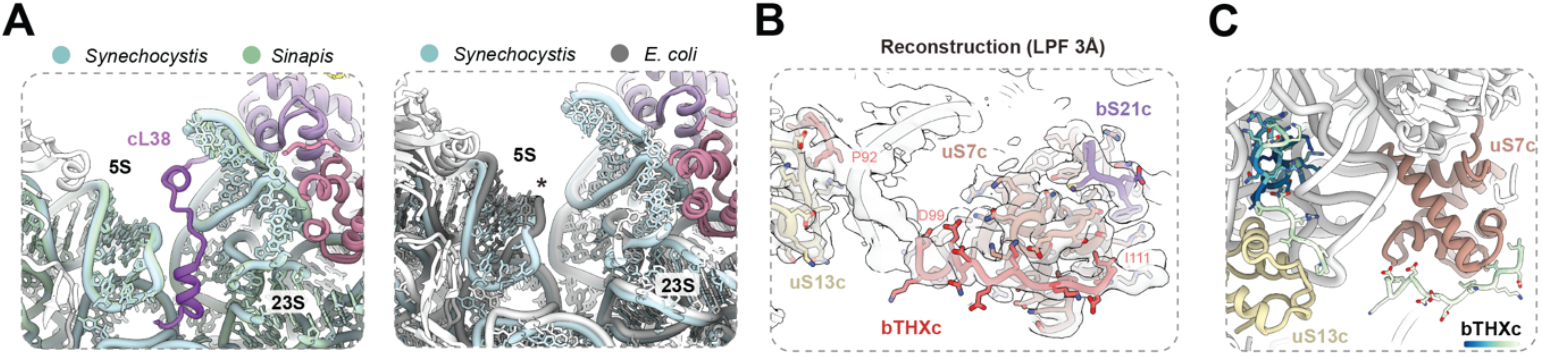
Additional data on PSRPs. **(A)** rRNA regions bound by PSRP6/cL38 in chloroplasts are structurally conserved in cyanobacteria and distinct from *E. coli*. Structural dissimilarity between the 5S hairpin D loop of *E. coli* and spinach proposed to be associated with the role of PSRP6/cL38 is indicated (asterisk). **(B)** Left: Details of the interaction between the C-terminal extension of bTHXc and uS7c. The cryo-EM reconstruction is shown low-pass filtered to 3Å due to higher B-factor of PSRP4 relative to its binding site on uS7c. **(C)** bTHXc in the chloroplast ribosome colored by residue-level conservation.

**Figure S7:**
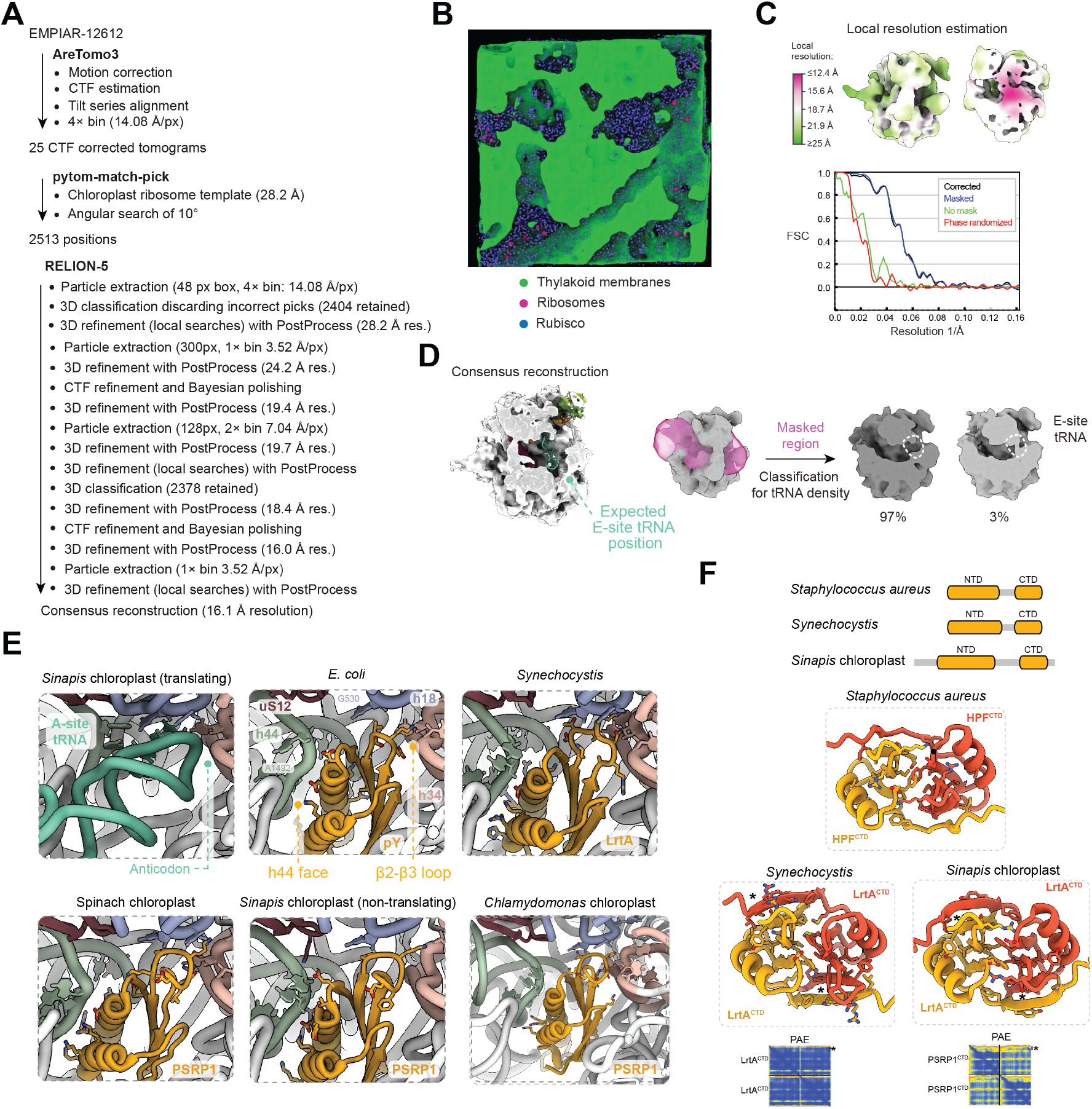
Additional data on the structural basis of ribosome hibernation in chloroplasts and cyanobacteria. **(A)** Cryo-ET data processing workflow leading to consensus in situ reconstruction of the spinach chloroplast ribosome. **(B)** Tomographic slice through isolated spinach chloroplast with annotation of thylakoid membranes, ribosomes and rubisco. **(C)** Fourier Shell Correlation (FSC) and local resolution estimates for consensus in situ reconstruction of the spinach chloroplast ribosome. **(D)** Three-dimensional classification of chloroplast ribosome states on the basis of density in the tRNA binding sites. **(E)** Structural comparison of the binding of N-terminal domains of HPF proteins to the ribosome A-site. Whereas the presence of interactions between α-helices of HPFs and 16S helix 44 (‘h44 face’) and between an HPF loop and the anticodon binding site are conserved, the specific residues that mediate contact are variable. Structures are shown for *E. coli* (PDB:6H4N) (49), spinach (PDB:6ERI) (4) and *Chlamydomonas* (PDB:28LU) (47). **(F)** Structural predictions of homodimerization of the C-terminal domains (CTDs) of HPF proteins generated using AlphaFold3. The predicted interactions for cyanobacterial and chloroplast HPF proteins involves β-sheet augmentation (asterisk).

**Table S1:**
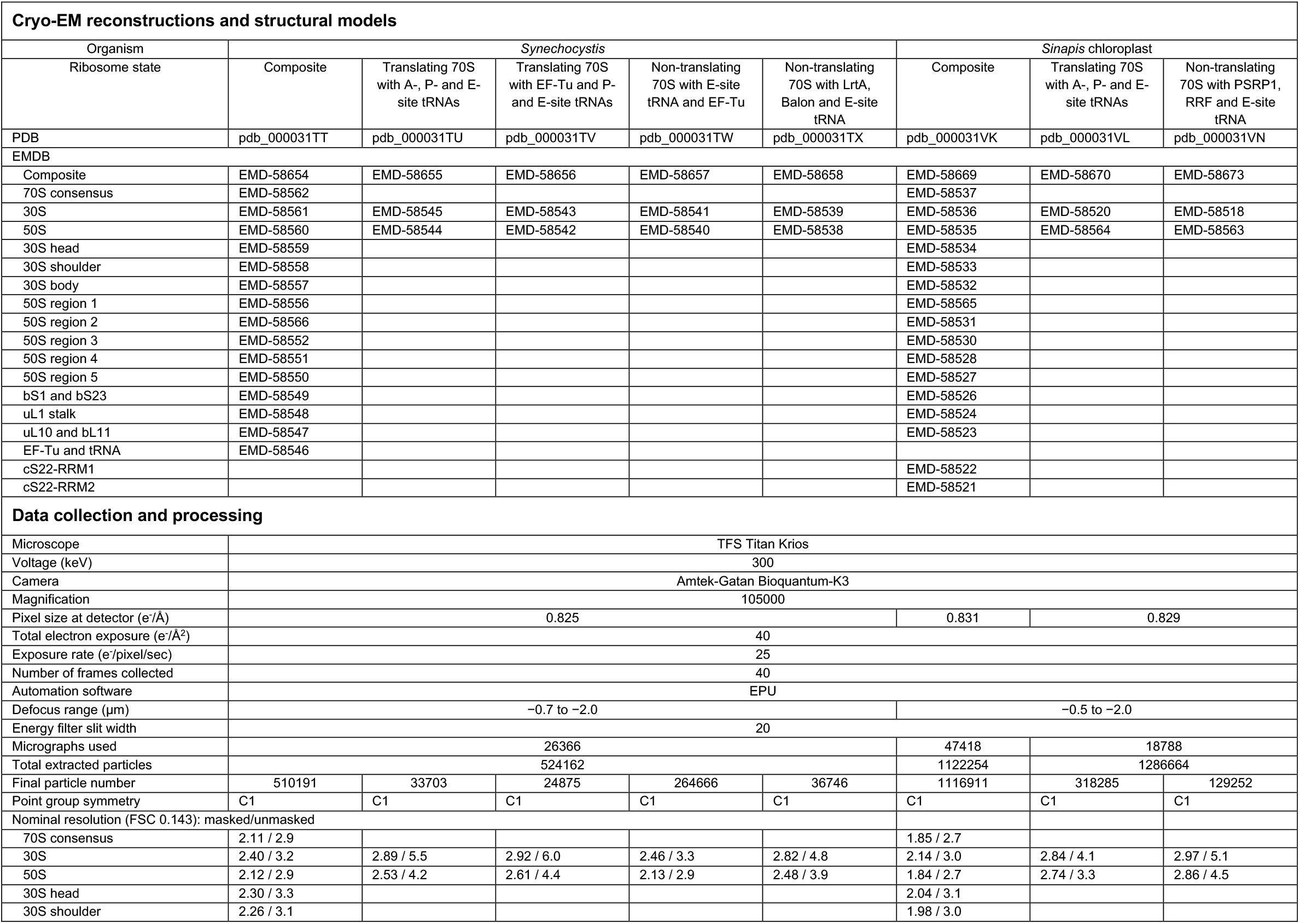
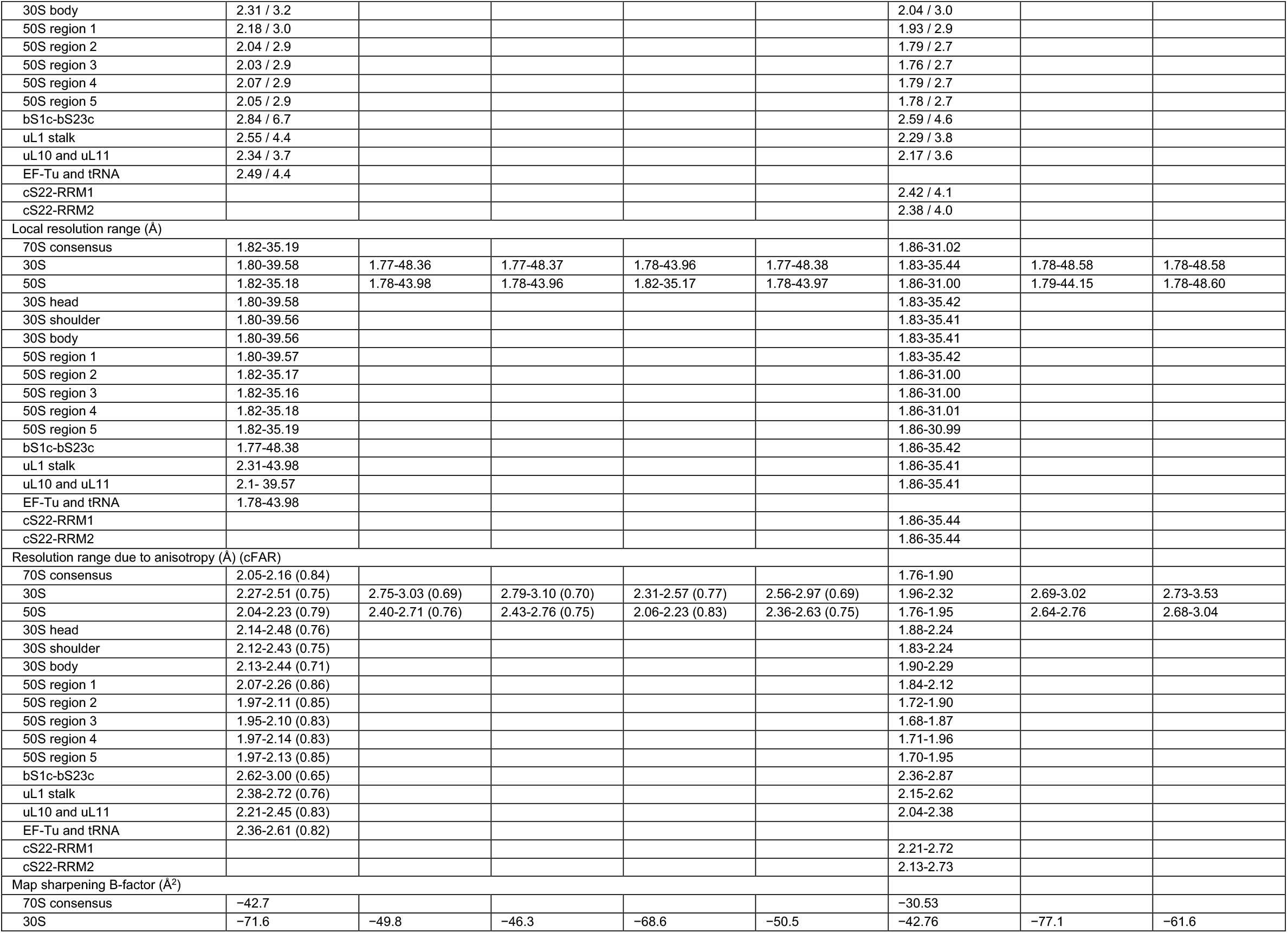
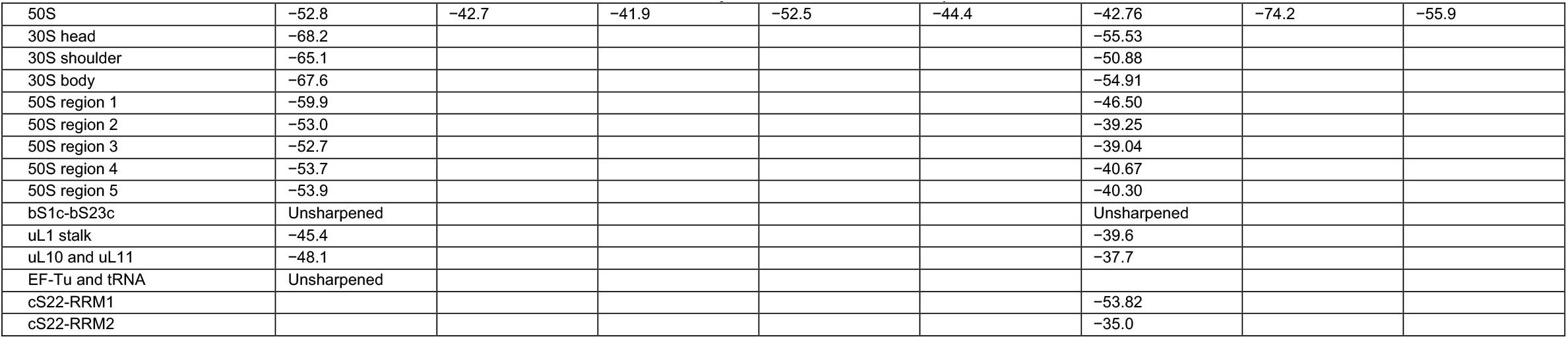
Cryo-EM data collection, refinement, and validation statistics.

| Cryo-EM reconstructions and structural models |  |  |  |  |  |  |  |  |
| --- | --- | --- | --- | --- | --- | --- | --- | --- |
| Organism | Synechocystis |  |  |  |  | Sinapis chloroplast |  |  |
| Ribosome state | Composite | Translating 70S with A-, P- and E-site tRNAs | Translating 70S with EF-Tu and P- and E-site tRNAs | Non-translating 70S with E-site tRNA and EF-Tu | Non-translating 70S with LrtA, Balon and E-site tRNA | Composite | Translating 70S with A-, P- and E-site tRNAs | Non-translating 70S with PSRP1, RRF and E-site tRNA |
| PDB | pdb_000031TT | pdb_000031TU | pdb_000031TV | pdb_000031TW | pdb_000031TX | pdb_000031VK | pdb_000031VL | pdb_000031VN |
| EMDB |  |  |  |  |  |  |  |  |
| Composite | EMD-58654 | EMD-58655 | EMD-58656 | EMD-58657 | EMD-58658 | EMD-58669 | EMD-58670 | EMD-58673 |
| 70S consensus | EMD-58562 |  |  |  |  | EMD-58537 |  |  |
| 30S | EMD-58561 | EMD-58545 | EMD-58543 | EMD-58541 | EMD-58539 | EMD-58536 | EMD-58520 | EMD-58518 |
| 50S | EMD-58560 | EMD-58544 | EMD-58542 | EMD-58540 | EMD-58538 | EMD-58535 | EMD-58564 | EMD-58563 |
| 30S head | EMD-58559 |  |  |  |  | EMD-58534 |  |  |
| 30S shoulder | EMD-58558 |  |  |  |  | EMD-58533 |  |  |
| 30S body | EMD-58557 |  |  |  |  | EMD-58532 |  |  |
| 50S region 1 | EMD-58556 |  |  |  |  | EMD-58565 |  |  |
| 50S region 2 | EMD-58566 |  |  |  |  | EMD-58531 |  |  |
| 50S region 3 | EMD-58552 |  |  |  |  | EMD-58530 |  |  |
| 50S region 4 | EMD-58551 |  |  |  |  | EMD-58528 |  |  |
| 50S region 5 | EMD-58550 |  |  |  |  | EMD-58527 |  |  |
| bS1 and bS23 | EMD-58549 |  |  |  |  | EMD-58526 |  |  |
| uL1 stalk | EMD-58548 |  |  |  |  | EMD-58524 |  |  |
| uL10 and bL11 | EMD-58547 |  |  |  |  | EMD-58523 |  |  |
| EF-Tu and tRNA | EMD-58546 |  |  |  |  |  |  |  |
| cS22-RRM1 |  |  |  |  |  | EMD-58522 |  |  |
| cS22-RRM2 |  |  |  |  |  | EMD-58521 |  |  |
| Data collection and processing |  |  |  |  |  |  |  |  |
| Microscope | TFS Titan Krios |  |  |  |  |  |  |  |
| Voltage (keV) | 300 |  |  |  |  |  |  |  |
| Camera | Amtek-Gatan Bioquantum-K3 |  |  |  |  |  |  |  |
| Magnification | 105000 |  |  |  |  |  |  |  |
| Pixel size at detector (e-/Å) | 0.825 |  |  |  |  | 0.831 | 0.829 |  |
| Total electron exposure (e-/Å²) | 40 |  |  |  |  |  |  |  |
| Exposure rate (e-/pixel/sec) | 25 |  |  |  |  |  |  |  |
| Number of frames collected | 40 |  |  |  |  |  |  |  |
| Automation software | EPU |  |  |  |  |  |  |  |
| Defocus range (µm) | -0.7 to -2.0 |  |  |  |  | -0.5 to -2.0 |  |  |
| Energy filter slit width | 20 |  |  |  |  |  |  |  |
| Micrographs used | 26366 |  |  |  |  | 47418 | 18788 |  |
| Total extracted particles | 524162 |  |  |  |  | 1122254 | 1286664 |  |
| Final particle number | 510191 | 33703 | 24875 | 264666 | 36746 | 1116911 | 318285 | 129252 |
| Point group symmetry | C1 | C1 | C1 | C1 | C1 | C1 | C1 | C1 |
| Nominal resolution (FSC 0.143): masked/unmasked |  |  |  |  |  |  |  |  |
| 70S consensus | 2.11 / 2.9 |  |  |  |  | 1.85 / 2.7 |  |  |
| 30S | 2.40 / 3.2 | 2.89 / 5.5 | 2.92 / 6.0 | 2.46 / 3.3 | 2.82 / 4.8 | 2.14 / 3.0 | 2.84 / 4.1 | 2.97 / 5.1 |
| 50S | 2.12 / 2.9 | 2.53 / 4.2 | 2.61 / 4.4 | 2.13 / 2.9 | 2.48 / 3.9 | 1.84 / 2.7 | 2.74 / 3.3 | 2.86 / 4.5 |
| 30S head | 2.30 / 3.3 |  |  |  |  | 2.04 / 3.1 |  |  |
| 30S shoulder | 2.26 / 3.1 |  |  |  |  | 1.98 / 3.0 |  |  |
| *Structures of the cyanobacterial and chloroplast ribosomes* |  |  |  |  |  |  |  |  |
| 30S body | 2.31 / 3.2 |  |  |  |  | 2.04 / 3.0 |  |  |
| 50S region 1 | 2.18 / 3.0 |  |  |  |  | 1.93 / 2.9 |  |  |
| 50S region 2 | 2.04 / 2.9 |  |  |  |  | 1.79 / 2.7 |  |  |
| 50S region 3 | 2.03 / 2.9 |  |  |  |  | 1.76 / 2.7 |  |  |
| 50S region 4 | 2.07 / 2.9 |  |  |  |  | 1.79 / 2.7 |  |  |
| 50S region 5 | 2.05 / 2.9 |  |  |  |  | 1.78 / 2.7 |  |  |
| bS1c-bS23c | 2.84 / 6.7 |  |  |  |  | 2.59 / 4.6 |  |  |
| uL1 stalk | 2.55 / 4.4 |  |  |  |  | 2.29 / 3.8 |  |  |
| uL10 and uL11 | 2.34 / 3.7 |  |  |  |  | 2.17 / 3.6 |  |  |
| EF-Tu and tRNA | 2.49 / 4.4 |  |  |  |  |  |  |  |
| cS22-RRM1 |  |  |  |  |  | 2.42 / 4.1 |  |  |
| cS22-RRM2 |  |  |  |  |  | 2.38 / 4.0 |  |  |
| Local resolution range (Å) |  |  |  |  |  |  |  |  |
| 70S consensus | 1.82-35.19 |  |  |  |  | 1.86-31.02 |  |  |
| 30S | 1.80-39.58 | 1.77-48.36 | 1.77-48.37 | 1.78-43.96 | 1.77-48.38 | 1.83-35.44 | 1.78-48.58 | 1.78-48.58 |
| 50S | 1.82-35.18 | 1.78-43.98 | 1.78-43.96 | 1.82-35.17 | 1.78-43.97 | 1.86-31.00 | 1.79-44.15 | 1.78-48.60 |
| 30S head | 1.80-39.58 |  |  |  |  | 1.83-35.42 |  |  |
| 30S shoulder | 1.80-39.56 |  |  |  |  | 1.83-35.41 |  |  |
| 30S body | 1.80-39.56 |  |  |  |  | 1.83-35.41 |  |  |
| 50S region 1 | 1.80-39.57 |  |  |  |  | 1.83-35.42 |  |  |
| 50S region 2 | 1.82-35.17 |  |  |  |  | 1.86-31.00 |  |  |
| 50S region 3 | 1.82-35.16 |  |  |  |  | 1.86-31.00 |  |  |
| 50S region 4 | 1.82-35.18 |  |  |  |  | 1.86-31.01 |  |  |
| 50S region 5 | 1.82-35.19 |  |  |  |  | 1.86-30.99 |  |  |
| bS1c-bS23c | 1.77-48.38 |  |  |  |  | 1.86-35.42 |  |  |
| uL1 stalk | 2.31-43.98 |  |  |  |  | 1.86-35.41 |  |  |
| uL10 and uL11 | 2.1- 39.57 |  |  |  |  | 1.86-35.41 |  |  |
| EF-Tu and tRNA | 1.78-43.98 |  |  |  |  |  |  |  |
| cS22-RRM1 |  |  |  |  |  | 1.86-35.44 |  |  |
| cS22-RRM2 |  |  |  |  |  | 1.86-35.44 |  |  |
| Resolution range due to anisotropy (Å) (cFAR) |  |  |  |  |  |  |  |  |
| 70S consensus | 2.05-2.16 (0.84) |  |  |  |  | 1.76-1.90 |  |  |
| 30S | 2.27-2.51 (0.75) | 2.75-3.03 (0.69) | 2.79-3.10 (0.70) | 2.31-2.57 (0.77) | 2.56-2.97 (0.69) | 1.96-2.32 | 2.69-3.02 | 2.73-3.53 |
| 50S | 2.04-2.23 (0.79) | 2.40-2.71 (0.76) | 2.43-2.76 (0.75) | 2.06-2.23 (0.83) | 2.36-2.63 (0.75) | 1.76-1.95 | 2.64-2.76 | 2.68-3.04 |
| 30S head | 2.14-2.48 (0.76) |  |  |  |  | 1.88-2.24 |  |  |
| 30S shoulder | 2.12-2.43 (0.75) |  |  |  |  | 1.83-2.24 |  |  |
| 30S body | 2.13-2.44 (0.71) |  |  |  |  | 1.90-2.29 |  |  |
| 50S region 1 | 2.07-2.26 (0.86) |  |  |  |  | 1.84-2.12 |  |  |
| 50S region 2 | 1.97-2.11 (0.85) |  |  |  |  | 1.72-1.90 |  |  |
| 50S region 3 | 1.95-2.10 (0.83) |  |  |  |  | 1.68-1.87 |  |  |
| 50S region 4 | 1.97-2.14 (0.83) |  |  |  |  | 1.71-1.96 |  |  |
| 50S region 5 | 1.97-2.13 (0.85) |  |  |  |  | 1.70-1.95 |  |  |
| bS1c-bS23c | 2.62-3.00 (0.65) |  |  |  |  | 2.36-2.87 |  |  |
| uL1 stalk | 2.38-2.72 (0.76) |  |  |  |  | 2.15-2.62 |  |  |
| uL10 and uL11 | 2.21-2.45 (0.83) |  |  |  |  | 2.04-2.38 |  |  |
| EF-Tu and tRNA | 2.36-2.61 (0.82) |  |  |  |  |  |  |  |
| cS22-RRM1 |  |  |  |  |  | 2.21-2.72 |  |  |
| cS22-RRM2 |  |  |  |  |  | 2.13-2.73 |  |  |
| Map sharpening B-factor (Å²) |  |  |  |  |  |  |  |  |
| 70S consensus | -42.7 |  |  |  |  | -30.53 |  |  |
| 30S | -71.6 | -49.8 | -46.3 | -68.6 | -50.5 | -42.76 | -77.1 | -61.6 |
| *Structures of the cyanobacterial and chloroplast ribosomes* |  |  |  |  |  |  |  |  |
| 50S | -52.8 | -42.7 | -41.9 | -52.5 | -44.4 | -42.76 | -74.2 | -55.9 |
| 30S head | -68.2 |  |  |  |  | -55.53 |  |  |
| 30S shoulder | -65.1 |  |  |  |  | -50.88 |  |  |
| 30S body | -67.6 |  |  |  |  | -54.91 |  |  |
| 50S region 1 | -59.9 |  |  |  |  | -46.50 |  |  |
| 50S region 2 | -53.0 |  |  |  |  | -39.25 |  |  |
| 50S region 3 | -52.7 |  |  |  |  | -39.04 |  |  |
| 50S region 4 | -53.7 |  |  |  |  | -40.67 |  |  |
| 50S region 5 | -53.9 |  |  |  |  | -40.30 |  |  |
| bS1c-bS23c | Unsharpened |  |  |  |  | Unsharpened |  |  |
| uL1 stalk | -45.4 |  |  |  |  | -39.6 |  |  |
| uL10 and uL11 | -48.1 |  |  |  |  | -37.7 |  |  |
| EF-Tu and tRNA | Unsharpened |  |  |  |  |  |  |  |
| cS22-RRM1 |  |  |  |  |  | -53.82 |  |  |
| cS22-RRM2 |  |  |  |  |  | -35.0 |  |  |

**Table S2:** Proteomic analysis of Sinapis chloroplast ribosome composition. Proteomics analysis of *Sinapis* chloroplast ribosome composition. Abundance is not shown for proteins in which only a single unique peptide was detected (*). Minor isoforms (‘isoform 2’) are shown for those that represent >10% of the total measured abundance.

| Protein | Isoform | NCBI/GenBank accession | Abundance (x10<sup>8</sup>) | Percent coverage |
| --- | --- | --- | --- | --- |
| uL1c | 1 | KAF8051660.1 | 17.0 | 56 |
| uL2c | 1 | YP_009730707.1 | 6.79 | 63 |
| uL3c | 1 | KAF8093973.1 | 8.44 | 63 |
| uL4c | 1 | KAF8116297.1 | 29.7 | 70 |
| uL5c | 1 | KAF8107115.1 | 10.8 | 81 |
| uL6c | 1 | KAF8106459.1 | 1.67 | 70 |
| bL9c | 1 | KAF8049991.1 | 17.0 | 64 |
| uL10c | 1 | KAF8109393.1 | 4.43 | 55 |
| uL11c | 1 | KAF8060378.1 | 15.8 | 63 |
| bL12c | 1 | KAF8049612.1 | 7.07 | 50 |
| bL12c | 2 | KAF8096717.1 | 4.81 | 48 |
| uL13c | 1 | KAF8103502.1 | * | 63 |
| uL14c | 1 | KAF8050889.1 | 9.23 | 83 |
| uL15c | 1 | KAF8096401.1 | 0.19 | 69 |
| uL15c | 2 | KAF8100894.1 | 0.06 | 69 |
| uL16c | 1 | KAF8100524.1 | 11.5 | 68 |
| bL17c | 1 | KAF8091157.1 | 4.09 | 33 |
| uL18c | 1 | KAF8048758.1 | 7.11 | 50 |
| bL19c | 1 | KAF8096239.1 | 0.95 | 55 |
| bL19c | 2 | KAF8098197.1 | 0.71 | 54 |
| bL20c | 1 | YP_009730689.1 | 0.78 | 41 |
| bL21c | 1 | KAF8053030.1 | 15.8 | 63 |
| uL22c | 1 | KAF8050892.1 | 0.11 | 38 |
| uL22c | 2 | KAF8044450.1 | 0.07 | 33 |
| uL23c | 1 | YP_009730708.1 | 0.16 | 48 |
| uL24c | 1 | KAF8092882.1 | 0.56 | 49 |
| uL24c | 2 | KAF8055305.1 | 0.50 | 49 |
| bL27c | 1 | KAF8050216.1 | 4.77 | 28 |
| bL28c | 1 | KAF8108829.1 | 0.95 | 58 |
| uL29c | 1 | KAF8089314.1 | 9.88 | 49 |
| bL31c | 1 | KAF8051977.1 | 4.00 | 56 |
| bL32c | 1 | YP_009730713.1 | 2.52 | 63 |
| bL33c | 1 | YP_009730687.1 | 0.75 | 68 |
| bL34c | 1 | KAF8075617.1 | 9.96 | 10 |
| bL35c | 1 | KAF8082910.1 | 11.2 | 11 |
| bL36c | 1 | YP_009730700.1 | 2.17 | 30 |
| cL37/PSRP5 | 1 | KAF8102088.1 | 0.35 | 45 |
| cL37/PSRP5 | 2 | KAF8049133.1 | 0.27 | 40 |
| cL37/PSRP5 | 3 | KAF8114734.1 | 0.07 | 35 |
| cL38/PSRP6 | 1 | KAF8085426.1 | 2.64 | 8 |
| bS1c | 1 | KAF8107216.1 | 3.85 | 53 |
| bS1c | 2 | KAF8105556.1 | 1.93 | 47 |
| uS2c | 1 | YP_009730656.1 | 2.00 | 61 |
| uS3c | 1 | KAF8048872.1 | 3.15 | 74 |
| uS4c | 1 | YP_009730669.1 | 5.70 | 59 |
| uS5c | 1 | KAF8108859.1 | 9.85 | 55 |
| bS6c | 1 | KAF8092289.1 | 3.70 | 34 |
| uS7c | 1 | YP_009730711.1 | 22.6 | 63 |
| uS8c | 1 | YP_009730701.1 | 2.91 | 80 |
| uS9c | 1 | KAF8111443.1 | 2.07 | 42 |
| uS10c | 1 | KAF8080879.1 | 4.41 | 22 |
| uS11c | 1 | YP_009730699.1 | 1.78 | 42 |
| uS12c | 1 | YP_009730646.1 | 3.11 | 52 |
| uS13c | 1 | KAF8118075.1 | 1.26 | 53 |
| uS14c | 1 | YP_009730665.1 | 6.46 | 53 |
| uS15c | 1 | KAF8047146.1 | 1.88 | 42 |
| bS16c | 1 | YP_009730649.1 | 0.92 | 21 |
| uS17c | 1 | KAF8096280.1 | 8.68 | 40 |
| uS17c | 2 | KAF8083859.1 | 3.54 | 39 |
| bS18c | 1 | YP_009730688.1 | 2.03 | 74 |
| uS19c | 1 | KAF8098587.1 | 0.28 | 29 |
| bS20c | 1 | KAF8086151.1 | 1.70 | 42 |
| bS20c | 2 | KAF8047656.1 | 0.62 | 46 |
| bS20c | 3 | KAF8100306.1 | 0.33 | 45 |
| bS21c | 1 | KAF8100931.1 | 0.11 | 20 |
| cS22/PSRP2 | 1 | KAF8107427.1 | 2.03 | 49 |
| cS22/PSRP2 | 2 | KAF8099760.1 | 0.28 | 43 |
| cS23/PSRP3 | 1 | KAF8094029.1 | 1.93 | 50 |
| bTHXc/PSRP4 | 1 | KAF8085305.1 | 4.65 | 40 |

**Table S5:**
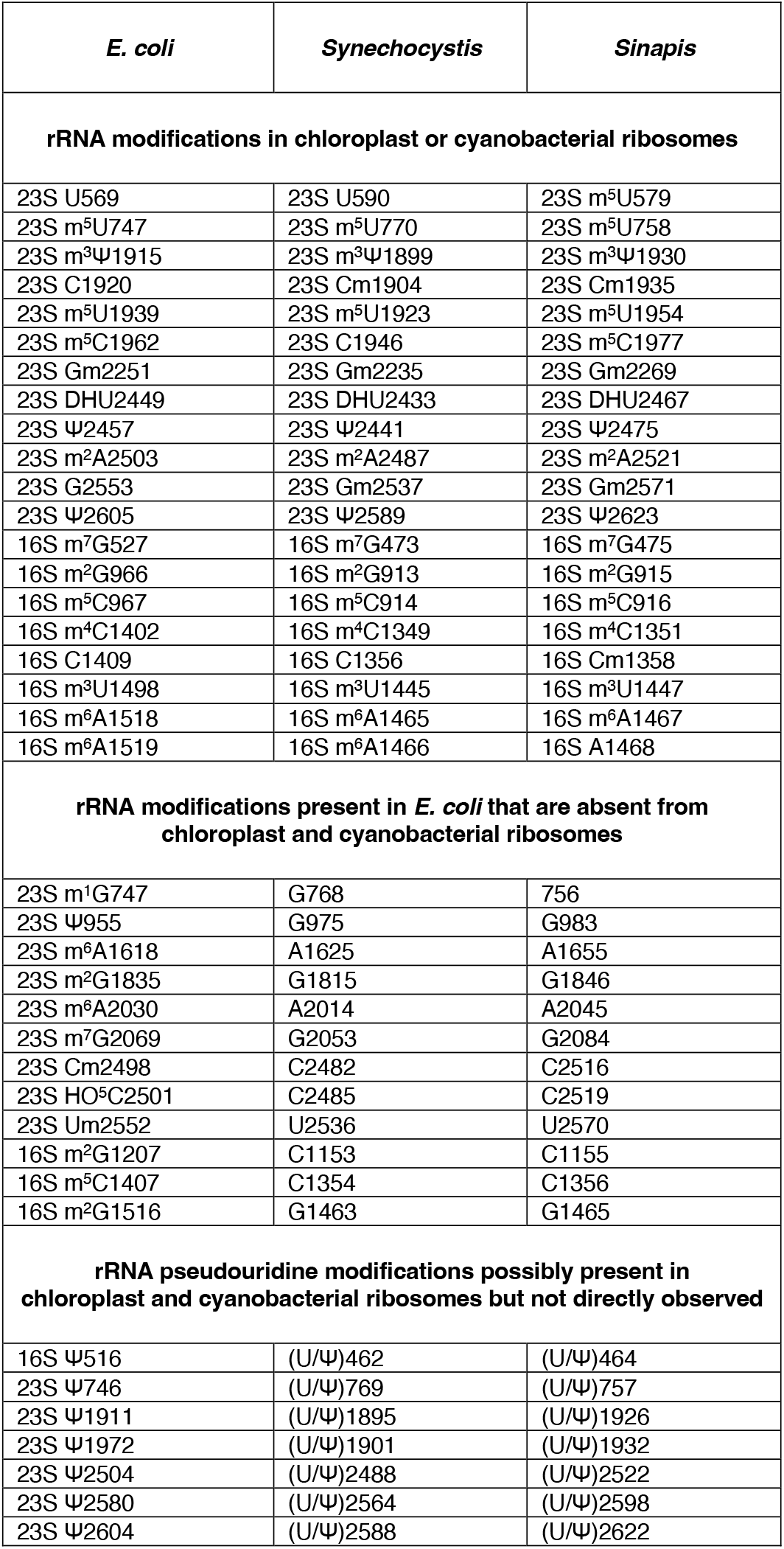
rRNA modifications in chloroplast or cyanobacterial ribosomes.

**Extended Data 1:**
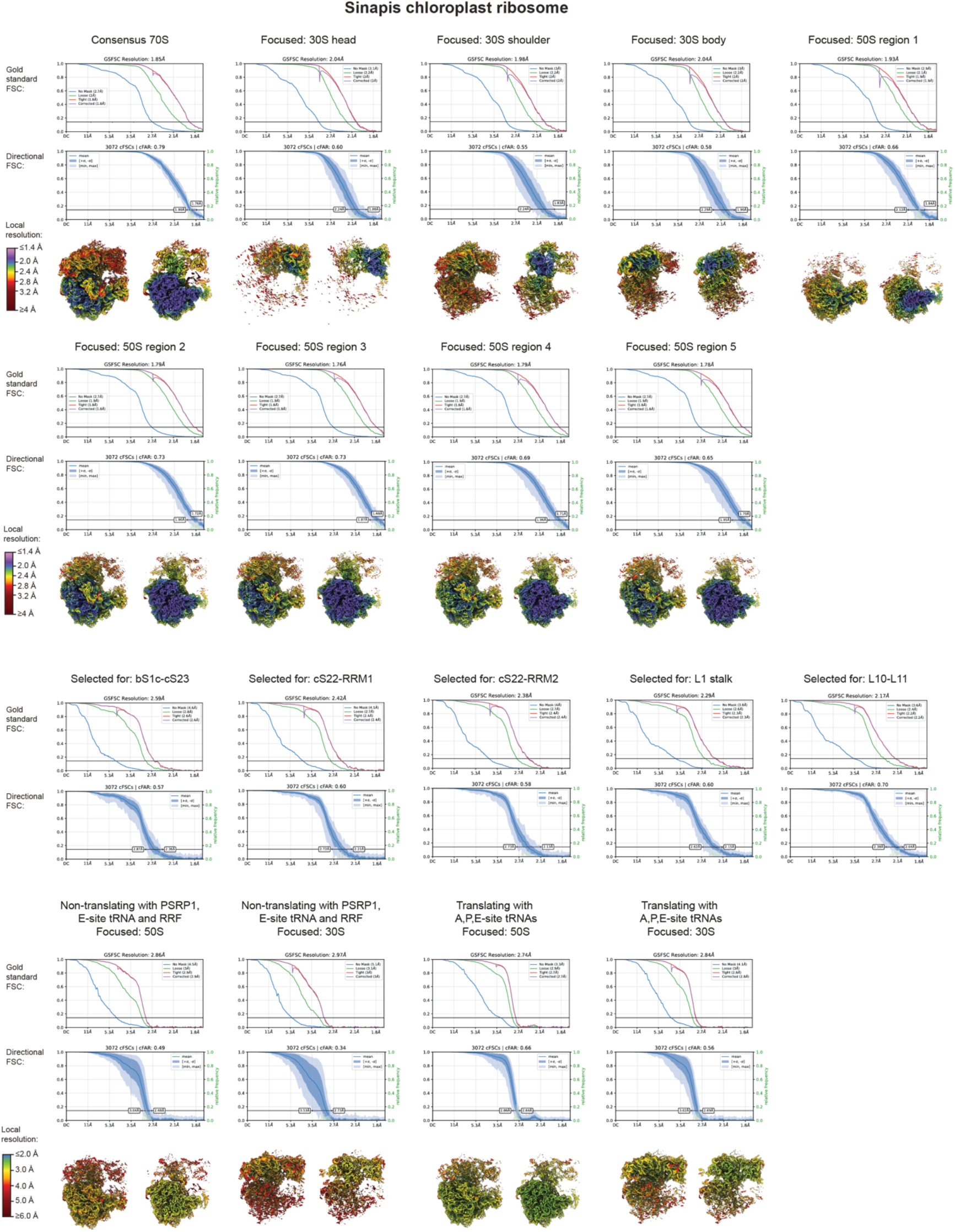
Resolution estimates for focused cryo-EM reconstructions of the *Sinapis* chloroplast ribosome.

**Extended Data 2:**
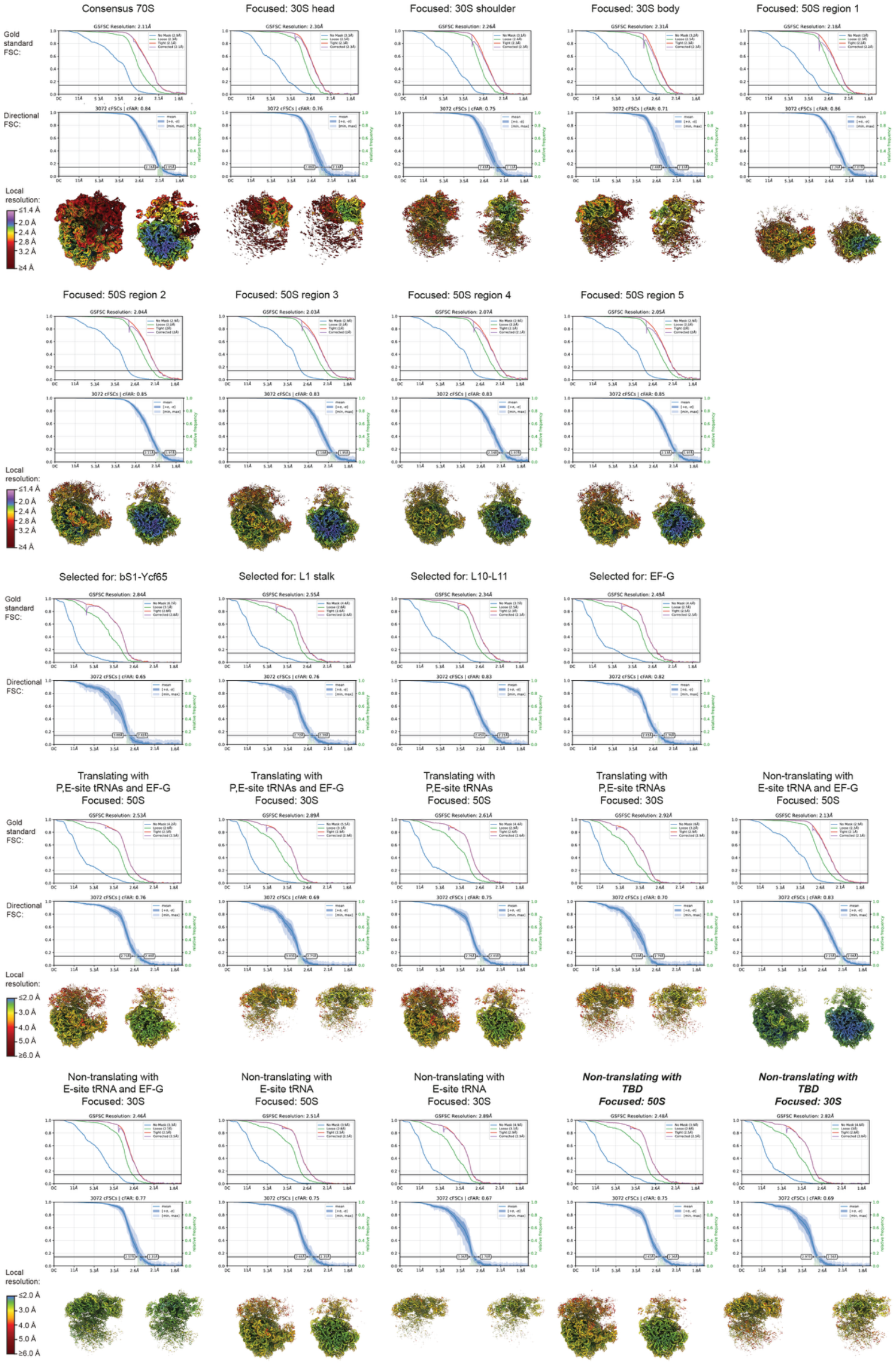
Resolution estimates for focused cryo-EM reconstructions of the *Synechocystis* ribosome.

